# 5-HT2C agonists and antagonists block different components of behavioral responses to potential, distal, and proximal threat in zebrafish

**DOI:** 10.1101/2020.10.04.324202

**Authors:** Rhayra Xavier do Carmo Silva, Bianca Gomes do Nascimento, Gabriela Cristini Vidal Gomes, Nadyme Assad Holanda da Silva, Jéssica Souza Pinheiro, Suianny Nayara da Silva Chaves, Ana Flávia Nogueira Pimentel, Bruna Patrícia Dutra Costa, Anderson Manoel Herculano, Monica Lima-Maximino, Caio Maximino

**Affiliations:** Laboratório de Neurofarmacologia Experimental – LNE, Universidade Federal do Pará, Belém/PA, Brazil; Laboratório de Neurociências e Comportamento “Frederico Guilherme Graeff” – LaNeC, Universidade Federal do Sul e Sudeste do Pará, Marabá/PA, Brazil; Laboratório de Neurofarmacologia e Biofísica – LaNeF, Universidade do Estado do Pará, Marabá/PA, Brazil

**Keywords:** 5-HT2C receptors, Anxiety, Fear, Responses to threat, Zebrafish

## Abstract

Serotonin (5-HT) receptors have been implicated in responses to aversive stimuli in mammals and fish, but its precise role is still unknown. Moreover, since at least seven families of 5-HT receptors exist in vertebrates, the role of specific receptors is still debated. Aversive stimuli can be classified as indicators of proximal, distal, or potential threat, initiating responses that are appropriate for each of these threat levels. Responses to potential threat usually involve cautious exploration and increased alertness, while responses to distal and proximal threat involve a fight-flight-freeze reaction. We exposed adult zebrafish to a conspecific alarm substance (CAS) and observed behavior during (distal threat) and after (potential threat) exposure, and treated with the 5-HT_2C_ receptor agonists MK-212 or WAY-161503 or with the antagonist RS-102221. The agonists blocked CAS-elicited defensive behavior (distal threat), but not post-exposure increases in defensive behavior (potential threat), suggesting inhibition of responses to distal threat. MK-212 blocked changes in freezing elicited by acute restraint stress, a model of proximal threat, while RS-102221 blocked changes in geotaxis elicited this stressor. We also found that RS-102221, a 5-HT_2C_ receptor antagonist, produced small effect on behavior during and after exposure to CAS.

**Preprint:** https://www.biorxiv.org/content/10.1101/2020.10.04.324202; **Data and scripts:** https://github.com/lanec-unifesspa/5-HT-CAS/tree/master/data/5HT2C

## 1. Introduction

Serotonergic mechanisms have been implicated in defensive behavior to distal, proximal and potential threat (Graeff, 2004). In current interpretations of the role of serotonin (5-HT) on neurobehavioral responses to threat, the neurotransmitter is released when threat is potential, increasing defensive behavior at this level (i.e., risk assessment/anxiety-like behavior), but inhibiting (non-adaptive) responses to distal and proximal threat (i.e., fight-flight-freeze/fear-like behavior)(Graeff, 2004). These effects are mediated by different structures of the neuroaxis, with more rostral structures (e.g., limbic forebrain) mediating responses to potential threat, and more caudal structures (e.g., periaqueductal gray/griseum centrale) mediating responses to distal and proximal threat (Graeff, 2004).

In zebrafish (*Danio rerio* Hamilton 1822), the alarm reaction has been proposed as a model to study defenses to distal threat (Maximino et al., 2019). The alarm reaction is a behavioral response to the release of an alarm substance (“Schreckstoff”) from epitelial club cells after damage (von Frisch, 1938), and involves strategies to avoid potential predators. Since the most likely situation for club cells to be damaged in the wild is predator attack, conspecific alarm substance (CAS) communicates to shoal-mates the potential presence of a predator, eliciting responses that avoid predator attack (Maximino et al., 2019). During CAS exposure, zebrafish display erratic swimming and bottom-dwelling, while after CAS exposure freezing is prominent (Lima-Maximino et al., 2020; Nathan et al., 2015). We suggested that the first component involves defenses to distal threat (“fear-like behavior”), while the second component involves a “return to normal” that is related to potential threat (“anxiety-like behavior”)(Lima-Maximino et al., 2020; Maximino et al., 2019).

5-HT has been implicated in fish behavioral responses both during and after exposure to CAS. In zebrafish, extracellular 5-HT levels were increased after CAS exposure (Maximino et al., 2014), an effect that can be related to decreased 5-HT uptake (Maximino et al., 2014) and/or decreased monoamine oxidase activity (Lima-Maximino et al., 2020; Quadros et al., 2018). In Crucian carp (*Carassius carassius*), exposure to CAS elicits increases in serotonergic activity in the brainstem and optic tectum (Höglund et al., 2005), structures which have been involved in responses to distal and proximal threat (do Carmo Silva et al., 2018a). However, this only happened when hiding material was unavailable in the tank. In Nile tilapia (*Oreochromis niloticus*), CAS did not increase serotonergic activity in the dorsomedial and dorsolateral telencephali (Silva et al., 2015), homologues of the frontotemporal amygdaloid nuclei and hippocampus, respectively (do Carmo Silva et al., 2018a). These results suggest that, during or after exposure, CAS increases serotonergic activity in regions associated with “quick-and-dirty” behavioral responses to distal threat (a “fight/flight/freeze” or “fear” system), but not in the telencephalic areas associated with cautious exploration/risk assessment (a “behavioral inhibition” or “anxiety” system).

Manipulations of the serotonergic system impact behavioral and neurovegetative responses to CAS. Treating zebrafish with acute fluoxetine, therefore increasing serotonergic activity, dose-dependently decreased behavior during exposure, but increased post-exposure freezing (Lima-Maximino et al., 2020). Blocking 5-HT receptors with metergoline, or depleting 5-HT with *para*-chlorophenylalanine, had no effect on behavior during exposure, but blocked the effects of CAS on post-exposure behavior. While we suggested that the serotonergic system is recruited *after* CAS exposure to inhibit fear-like responses and promote a cautious “return to normal” (Lima-Maximino et al., 2020), results from Crucian carp (Höglund et al., 2005) open the possibility that inescapability is the variable that is involved in this activation of the serotonergic system.

The role of serotonin receptors from the 5-HT_2_ family in CAS-elicited behavioral adjustments has also been investigated. Zebrafish has been shown to possess two copies of the 5-HT_2A_ receptor, one copy of the 5-HT_2B_ receptor, and two copies of the 5-HT_2C_ receptor (Sourbron et al., 2016). In Nile tilapia, mianserin, a 5-HT_2A_ and 5-HT_2C_ receptor antagonist that also blocks α_2_-adrenoceptors, blocked active components of the alarm reaction (dashing, bristling of dorsal fin spines), but not freezing, during exposure (Barreto, 2012). In zebrafish, methysergide (an antagonist at 5-HT_2A_, 2B, and 2C receptors, and a 5-HT_1A_ receptor agonist) increased freezing and bottom-dwelling during and after exposure at a high dose (92.79 mg/kg), but not at lower doses (Nathan et al., 2015). These results point to an inhibitory role of the 5-HT_2A_ and 5-HT_2C_ receptors on CAS-elicited fear-like responses. However, since Nathan et al. (2015) observed these effects consistently during a long session, in which behavior is expected to change as CAS concentrations decrease (Mathuru et al., 2012)(i.e., similar to the shift from erratic swimming to freezing that is observed when animals are observed after exposure in a CAS-free context; Lima-Maximino et al., 2020), and since serotonergic drugs can produce opposite effects on behavior during and after CAS exposure (Lima-Maximino et al., 2020), from these results it is not possible to understand whether 5-HT_2_ receptors participate in both responses. In rodents, for example, 5-HT_2_ receptors appear to inhibit fear and facilitate anxiety at different levels of the neuroaxis (e.g. Graeff et al., 1986; Coimbra and Brandão, 1997; Castilho and Brandão, 2001; Castilho et al., 2002; Alves et al., 2004; Cornélio and Nunes-de-Souza, 2007; Macedo et al., 2007; Nunes-de-Souza et al., 2008; de Paula and Leite-Panissi, 2016), while in zebrafish these receptors appear to inhibit both fear and anxiety, although it is currently unknown which brain regions participate in each effect.

In this work, we tested whether 5-HT_2C_ receptors participate in responses to CAS during (distal threat) or after exposure (potential threat) and to acute restraint stress (ARS). ARS has been applied in zebrafish to elicit strong stress responses, including activation of the hypothalamus-pituitary-interrenal (HPI) axis and associated behavioral responses (Assad et al., 2020; Ghisleni et al., 2012; Piato et al., 2011). From the point of view of the predatory imminence continuum theory, ARS represents proximal threat (Perusini and Fanselow, 2015). Thus, if 5-HT_2C_ receptors inhibit aversively motivated behavior in zebrafish regardless of threat distance, its activation would inhibit behavioral responses at these three contexts. If, as in rodents, 5-HT_2C_ receptors act at different levels of threat distance to either activate or inhibit defensive responses, then 5-HT_2C_ agonists will not produce the same effect in each of these contexts.

The aim of the present work was to test the hypothesis that activation of 5-HT_2C_ receptors would block CAS-elicited defensive reactions during exposure, but not after exposure; moreover, we proposed that 5-HT_2C_ receptor antagonists would affect post-exposure behavior. We also hypothesized that 5-HT_2C_ agonists would not alter the anxiogenic-like effects of acute restraint stress (ARS), a model for proximal threat. We found that 5-HT_2C_ agonists were able to block elements of the alarm reaction, but did not affect post-exposure behavior, nor the full anxiogenic-like effects of ARS, while a 5-HT_2C_ receptor antagonist partially blocked the alarm reaction and more strongly affected post-exposure behavior. This manuscript is a complete report of all the studies performed to test the effect of 5-HT_2C_ agonists and antagonists on zebrafish defensive behavior to distal threat. We report how the sample size was determined, all data exclusions (if any), all manipulations, and all measures in the study.

## 2. Materials and methods

### 2.1. Animals and housing

Adult (>4 month-old; standard length = 23.0 ± 3.2 mm) zebrafish (*Danio rerio*) from the longfin phenotype (n = 250) were used in the present experiments. The populations used are expected to better represent the natural populations in the wild, due to its heterogeneous genetic background (Parra et al., 2009; Speedie and Gerlai, 2008). Animals were bought from a commercial vendor (Belém/PA) and collectively maintained in 40 L tanks for at least two weeks before the onset of experiments. The animals were fed daily with fish flakes. The tanks were kept at constant temperature (28 °C), oxygenation, light cycle (14:10 LD photoperiod) and a pH of 7.0-8.0, according to standards of care for zebrafish (Lawrence, 2007). Animals were used for only one experiment to reduce interference from apparatus exposure. Potential suffering of animals was minimized by controlling for the aforementioned environmental variables. Furthermore, in the all experiments the animals used were handled, anesthetized and sacrificed according to the norms of the Brazilian Guideline for the Care and Use of Animals for Scientific and Didactical Purposes (Conselho Nacional de Controle de Experimentação Animal - CONCEA, 2017). Since, in all cases, animals were individually stressed and/or individually treated, experimental unit was a single animal, and therefore all sample sizes refer to number of animals. The experimental protocols were approved by UEPA’s IACUC under protocol 06/18.

### 2.2. Drugs and treatments

The 5-HT_2C_ receptor agonists MK-212 (2-Chloro-6-(1-piperazinyl)pyrazine, CAS #64022-27-1) and WAY-161503 (8,9-dichloro-2,3,4,4a-tetrahydro-1H-pyrazino[1,2-a]quinoxalin-5(6H)-one; CAS #75704-24-4) were bought from Sigma-Aldrich (St Louis, USA) on 2018, and dissolved in Cortland’s salt solution (NaCl 124.1 mM, KCl 5.1 mM, Na_2_HPO_4_ 2.9 mM, MgSO_4_ 1.9 mM, CaCl_2_ 1.4 mM, NaHCO_3_ 11.9 mM, Polyvinylpyrrolidone 4%, 1,000 USP units Heparin; Wolf, 1963) and in 1% dimethyl sulfoxide (DMSO), respectively. The 5-HT_2C_ receptor antagonist RS-102221 (8-[5-(2,4-Dimethoxy-5-(4-trifluoromethylphenylsulphonamido)phenyl-5-oxopentyl]-1,3,8-triazaspiro[4.5]decane-2,4-dione; CAS #185376-97-0) was bought from Sigma-Aldrich (St Louis, USA) in 2018, and dissolved in 1% DMSO. While affinities for zebrafish 5-HT_2C_-like receptors have not been established, WAY-161503 has been reported to displace DOI from human 5-HT_2C_ receptors with a *K_i_* of 3.3 nM (6-fold selectivity over human 5-HT_2A_ receptors and 20-fold over human 5-HT_2B_ receptors)(Rosenzweig-Lipson et al., 2006). MK-212 has been shown to be less selective at recombinant human receptors, with a *K_i_* of 7.01 nM at 5-HT_2C_ receptors (vs. 5.99 nM and 6.21 nM at 5-HT_2A_ and 5-HT_2B_ receptors, respectively)(Knight et al., 2004). Finally, RS-102221 has been shown to displace mesulergine from human 5-HT2C receptor with a *pKi* of 8.4 nM (over 100-fold selectivity over human 5-HT_2A_ and 5-HT_2B_ receptors) (Bonhaus et al., 1997).

For Experiment 1, animals were injected intraperitoneally either with MK-212 (1 mg/kg and 2 mg/kg, doses which increase anxiety-like behavior in the rat elevated plus-maze; (de Mello Cruz et al., 2005)) or with the vehicle solution (Cortland’s salt solution); WAY-161503 (1 mg/kg, a dose which produces anxiogenic-like effects in the rat elevated plus-maze: Gomes et al., 2010) or with the vehicle solution (DMSO); or RS-102221 (2 mg/kg, a dose that reduces anxiety in the mouse light/dark test; (Kuznetsova et al., 2006)). For Experiment 2, animals were injected intraperitoneally with MK-212 (2 mg/kg) or vehicle (Cortland’s salt solution). Injections were made according to the protocol proposed by Kinkel et al. (2010); briefly, animals were cold-anesthetized and transferred to a sponge-based surgical bed, in which injection was made. Injections were made using a microsyringe (Hamilton® 701N syringe, needle size 26 gauge at cone tip), with total volumes of injection of 5 μL.

### 2.3. Experiment 1: Effects of 5-HT_2C_ receptor agonists and antagonists on alarm reaction and post-exposure behavior

#### 2.3.1. Experimental design

To verify the effects of phasic activation of 5-HT_2C_ receptors on the zebrafish alarm reaction, animals were pre-treated with either receptor agonists and exposed to CAS in a sequential design, with a “washout” period in between tests (Figure 1A). For the exposure stage, each animal was transferred individually to a container (2 L) where after 3 minutes of acclimatization, it was carefully exposed to 7 ml of alarm substance (CAS), extracted using a standardized protocol (do Carmo Silva et al., 2018b). As negative control, a group with the same amount of animals was exposed to the same volume of distilled water, according to the protocol of Lima-Maximino et al. (2020). The animals remained exposed for 6 minutes during which their behavior was recorded using a video camera positioned in front of the aquarium. Then, to verify the residual effects of exposure to the alarm substance, the animals were transferred to the apparatus of the novel tank test, a transparent glass aquarium filled with 5 L of mineral water where the animal can freely explore the space for a period 6 minutes during which their behavior was recorded, following the protocol described in Lima-Maximino et al., (2020). All stages of the experiment were performed under constant white Gaussian noise, producing an average of 58 dB above the tank. Light levels above the tanks were measured using a handheld light meter, and ranged from 251 to 280 lumens (coefficient of variation = 3.399% between subjects).

**Figure 1.**
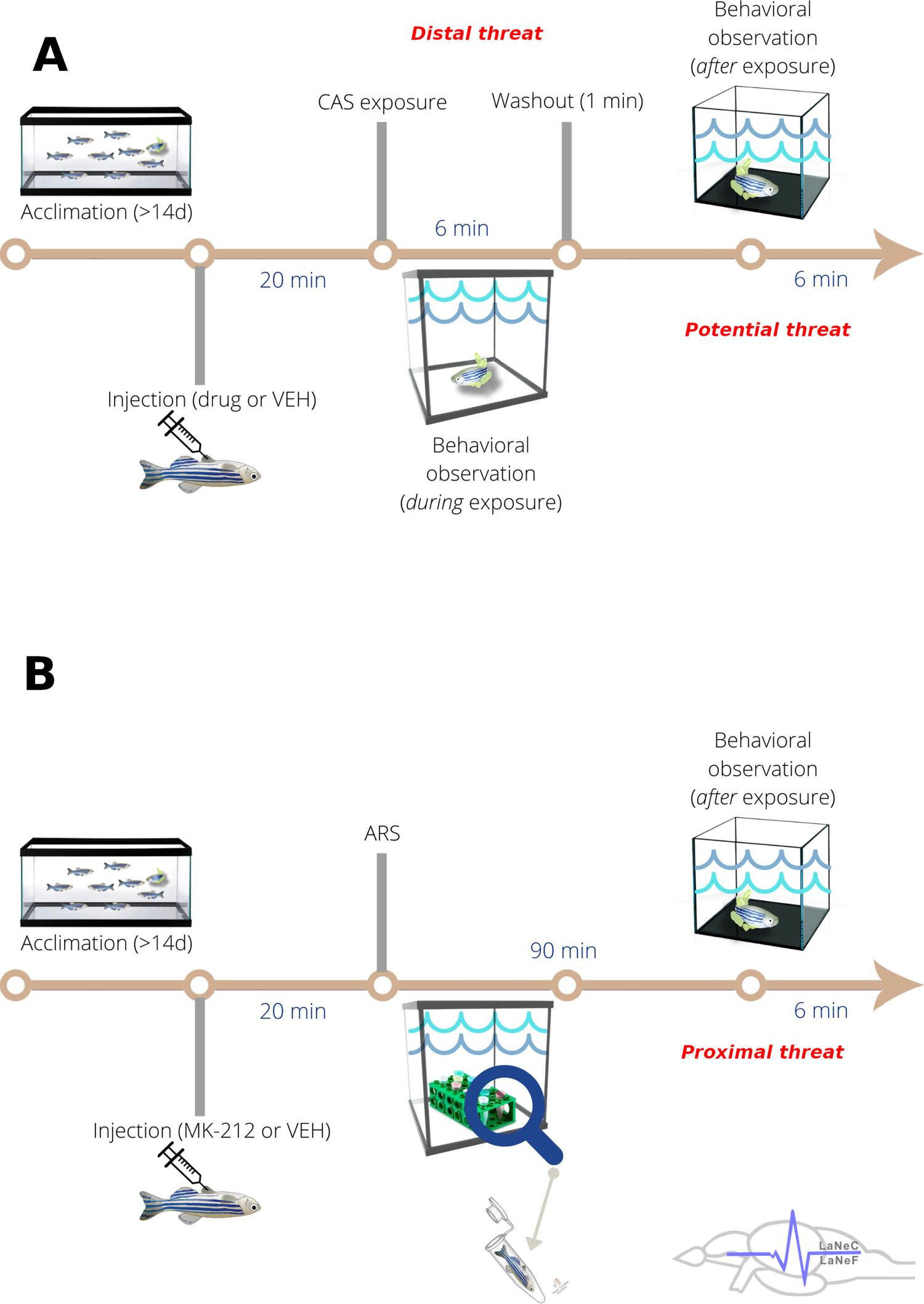
Experimental designs for (A) experiment 1 (CAS-elicited behavioral responses [distal threat] and post-exposure behavior [potential threat]) and (B) experiment 2 (ARS-elicited behavioral responses [proximal threat]). *Abbreviations:* ARS – acute restraint stress; CAS – conspecific alarm substance; VEH – vehicle

Animals were randomly allocated to groups using a random number generator (http://www.jerrydallal.com/random/random_block_size_r.htm), with each subject randomized to a single treatment using random permuted blocks. For each experiment, animals were treated and tested in the order of allocation (i.e., randomly). In all experiments, experimenters and data analysts were blinded to drugs and treatment by using coded vials (with the same code used for randomization); blinding was removed only after data analysis. Experiments were always run between 08:00AM and 02:00 PM. After experiments, animals were sacrificed by prolonged bath in ice-cold water (< 12 °C), followed by spinal transection (Matthews and Varga, 2012).

#### 2.3.2. Sample size determination

To determine sample size, we incorporated information from control groups derived from previously published experiments on zebrafish alarm substances observing bottom-dwelling (Lima-Maximino et al., 2020; Quadros et al., 2016; Speedie and Gerlai, 2008) and a series of four small experiments on the effects of CAS on behavior during exposure (https://github.com/lanec-unifesspa/5-HT-CAS/tree/master/data/behavioral/metanalysis), following the RePAIR approach (Bonapersona et al., 2019). Sample sizes, means, and standard deviations for the primary endpoint “Time on bottom” were used to produce a prior distribution on the RePAIR script (https://utrecht-university.shinyapps.io/repair/). Final parameters of the distribution were μ = 197.149, σ² = 6100.609, and weighted *N* = 120.10. The parameters of this distribution were then used to calculate sample size, based on an effect size of *d* = 0.5 (equivalent to that used to calculate sample sizes in Lima-Maximino et al., 2020) and *a priori* power of 0.8, with one-tailed tests with error probability α = 0.05. With these parameters, the final number of animals was 10 animals in the control group and 21 in each experimental group, reaching a prospective power of 92.34%.

#### 2.3.3. Alarm substance extraction

A group of zebrafish was used as donor animals for the extraction of the conspecific alarm substance (CAS). CAS extraction procedure was performed on each animal individually as described by do Carmo Silva et al. (2018b). First, the donor animal was cold anesthetized and transferred to a Petri dish, where the excess water from its body was removed with a paper towel. Then the animal was decapitated with a surgical scalpel and the excess blood from the sectioned region was removed with a swab. Subsequently, the animal’s bodies were transferred to another Petri dish where 15 superficial cuts were made in the epidermis of animals (medial-ventral region) and 10 ml of distilled water were added to wash the cuts. After washing, the animal’s bodies were removed from the Petri dish, and with the aid of a Pasteur pipette, the fish scales and other impurities were removed from the solution that was stored in a conical tube and preserved on ice. The same extraction procedure was performed for all donated animals.

#### 2.3.4. Observation apparatuses

To assess the effects of drugs on the alarm reaction, animals were transferred to a 12 cm x 12 cm x 14 cm glass tank, filled with 1.5 L tank water, and allowed to swim freely for a 3 min acclimation period. After this period, 7 mL CAS was added to the tank, from above, and filming started. The video camera (Sony® DCR-DVD610) was positioned in the front of the tank, therefore allowing observation and tracking of vertical distribution. Animals were allowed to freely explore this tank for 6 min before being transferred to a “washout” tank, which contained 500 mL tank water. Animals were kept in this tank for 1 min, removing any residues of CAS before transference to the second apparatus. This second apparatus was a 25 cm x 24 cm x 20 cm glass tank, filled with 5 L tank water; animals were allowed free exploration of this tank for 6 min, during which behavior was filmed from the front of the tank. Tanks for both stages were differently shaped to increase the novelty of the second environment, a variable that is important to induce an anxiety-like “diving” response in animals not exposed to CAS (Bencan et al., 2009).

### 2.4. Experiment 2: Effects of MK-212 or RS-102221 on acute restraint stress-elicited behavior

#### 2.4.1. Experimental design

In this experiment, we focus on assessing the roles of 5-HT_2C_ receptors on anxiogenic-like effects of acute restraint stress (ARS). In order to do this, we evaluate the behavior of the animals in the novel tank test after being subjected to a 90-min section of restraint stress (Figure 1B). After drug (MK-212 or RS-102221) or vehicle (Cortland’s salt solution) injection and anesthesia, each animal was transferred individually to a 2 mL microtube (Eppendorf®) and placed on a plastic microtube rack inside an aquarium with continuous oxygen supply. The microtubes had small holes at both ends to allow free circulation of water inside the tube and to prevent fish from moving around, according to the protocol of Piato et al. (2011). A control group was maintained in a similar tank, but without restraint stress. Animals remained in these conditions for 90 min., sufficient to induce changes in telencephalon neurochemistry (Assad et al., 2020) and marked anxiety-like behavior (Assad et al., 2020; Ghisleni et al., 2012; Piato et al., 2011). Animals in the control groups were subjected to all manipulations with the exception of restraint. After stress, the animals were transferred to the apparatus of the novel tank test, a transparent glass aquarium filled with 5 L of mineral water where the animal can freely explore the space for a period 6 minutes during which their behavior was recorded, following the protocol described in Lima-Maximino et al., (2020). All stages of the experiment were performed under constant white Gaussian noise, producing an average of 58 dB above the tank. Light levels above the tanks were measured using a handheld light meter, and ranged from 254 to 276 lumens (coefficient of variation = 3.401% between subjects). Random allocation was made as described above. In all experiments, experimenters and data analysts were blinded to drugs by using coded vials (with the same code used for randomization); blinding was removed only after data analysis. Experimenters were not blinded to treatment, but data analysts were blinded to both treatment and drug. Experiments were always run between 08:00AM and 02:00 PM. After experiments, animals were sacrificed by prolonged bath in ice-cold water (< 12 °C), followed by spinal transection (Matthews and Varga, 2012).

#### 2.4.2. Sample size determination

Sample sizes were based on previous experiments with the behavioral effects of ARS (Assad et al., 2020), using the same parameters as the CAS sample size determination, with the exception of effect size (which was defined as *d* = 4.56263, based on Assad et al., 2020).

#### 2.4.3. Observation apparatus

To analyze the behavioral effects of treatment (ARS exposure) and drug, the same apparatus used in the second stage of the previous experiment (5-L transparent glass tank) was used.

### 2.5. Behavioral endpoints

Video files for each experiment were stored and later analyzed using automated video tracking (TheRealFishTracker; http://www.dgp.toronto.edu/~mccrae/projects/FishTracker/). The following variables were extracted:

- Time spent on bottom third of the tank (s)[Primary outcome]
- Time spent on top third of the tank (s)[Secondary outcome]
- Erratic swimming, measured as absolute turn angle [Secondary outcome]
- Freezing (s), measured as time spent in a speed lower than 0.5 cm/s [Secondary outcome]
- Swimming speed (cm/s) [Secondary outcome]

### 2.6. Quality control

**Exclusion criteria:** Animals were to be excluded if showing signs of poor health or distress before beginning experiments, using the Zebrafish Health and Welfare Glossary (https://wiki.zfin.org/display/ZHWG/Zebrafish+Health+and+Welfare+Glossary+Home; Goodwin et al., 2016). Moreover, animals could be excluded from analysis based on statistical detection as outlier (see 2.7, “Statistical analysis”, below).

**Behavioral data**: Quality control of samples was maintained by periodic assessment of water quality and health parameters. All experimenters were trained in the behavioral methods before experiments; training included observation of all experiments by a PI (CM or MGL) on at least two occasions. After these observations, each trainee performed two mock experiments, on a single subject each, while being observed by the PI. All protocols were reviewed by all PIs, and are publicly available. Behavioral records were reviewed by at least one PI for administration/scoring accuracy, in order to ensure adherence to protocols and consistency across tests.

### 2.7. Statistical analysis

Data were analyzed using two-way analysis of variance (drug x exposure to stressor) with sequential sum of squares (type I), followed by Tukey’s post-tests when p <0.05; since multiple comparisons were made in post-tests, p-values were calculated using Tukey’s method. Data analysis, table organization, and result graphs were performed using R version 3.6.3 (2020-02-29). Effect sizes for ANOVA effects are shown as ω²; effect sizes for post-hoc tests were shown as Cohen’s d. Outliers were removed based on median absolute differences (MADs), using time on bottom as the main endpoint; values were removed when they were higher or lower than 3 MADs around the median (Leys et al., 2013), and the number of outliers was reported in the results.

### 2.8. Sequence alignment

Zebrafish have two copies of the 5-HT2C receptor gene, *htr2cl1* [ENSDARG00000018228] and *htr2cl2* [ENSDARG00000013210] (Sourbron et al., 2016); currently, only the expression of *htr2cl1* is known, and studies were made in larvae (Schneider et al., 2012). Moreover, receptor binding and drug affinity is currently unknown. Mutation and *in silico* studies suggest that the 5-HT_2C_ receptor residues D3.32, S3.36, and Y7.43 are crucial to ligand binding (Canal et al., 2011; Córdova-Sintjago et al., 2012). We looked for the conservation of these residues across zebrafish 5-HT_2C-_like receptors by aligning protein sequences from zebrafish receptors (XP_017212271.1 and XP_001339040.4) with the murine (NP_032338.3) and human (NP_001243689.2) 5-HT_2C_ receptors in Clustal Omega (https://www.ebi.ac.uk/Tools/msa/clustalo/). Resulting alignments were visually inspected for conserved residues at the positions that are related to ligand affinity.

### 2.9 Open science practices

The studies presented here were not formally pre-registered. A preprint version was posted to bioRxiv (https://www.biorxiv.org/content/10.1101/2020.10.04.324202v2. Data and analysis scripts can be found on a Github repository (https://github.com/lanec-unifesspa/5-HT-CAS/tree/master/data/5HT2C).

## 3. Results

### 3.1. Effects of MK-212 on alarm reaction and post-exposure behavior

#### 3.1.1. During exposure

One animal from the control + 0 mg/kg group was detected as outlier and removed. Full results for ANOVAs can be found on Table 1. Post-hoc tests found that CAS did not alter time on top (p = 0.94, d = 0.3, non-treated controls vs. non-treated CAS), but MK-212 (1 mg/kg) increased it in non-exposed animals (*1 mg/kg:* p = 0.024, d = −1.05, non-treated controls vs. 1 mg/kg controls; *2 mg/kg:* p = 0.994, d = −0.18, non-treated controls vs. 2 mg/kg controls; Figure 2A).

**Figure 2.**
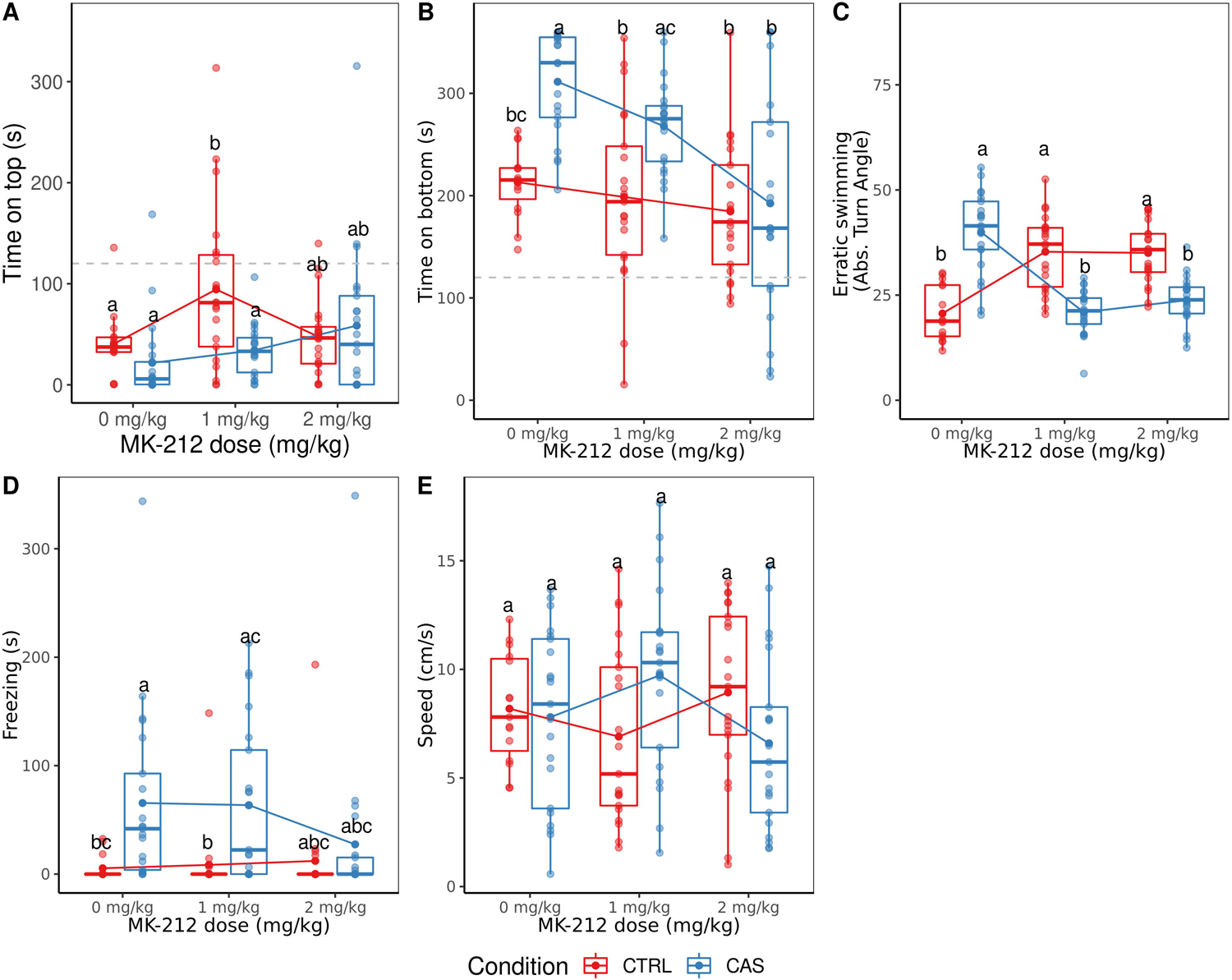
Effects of MK-212 on behavior <u>during</u> CAS exposure. (A) Time spent on top third of the tank. (B) Time spent on bottom third of the tank. (C) Erratic swimming. (D) Total time spent freezing. (E) Swimming speed. Different letters represent statistical differences at the p < 0.05 level; similar letters indicate lack of statistically significant differences. Data are presented as individual data points (dots) superimposed over the median ± interquartile ranges. Dashed lines on panels A and B represent change levels. Dots connected by lines represent group means. CTRL = controls (water-exposed animals); CAS = conspecific alarm substance. Final sample sizes: CTRL + VEH: n = 15 animals; CTRL + 1 mg/kg MK-212: n = 21 animals; CTRL + 2 mg/kg MK-212: n = 21 animals; CAS + VEH: n = 21 animals; CAS + 1 mg/kg MK-212: n = 21 animals; CAS + 2 mg/kg MK-212: n = 21 animals.

**Table 1.** Full ANOVA results for effects of MK-212 on behavioral endpoints during CAS exposure.

| Context | Drug | Endpoint | Effect<br>[df, df2] | F | p-value | Effect size<br>($\omega^2$) |
| --- | --- | --- | --- | --- | --- | --- |
| During exposure | MK-212 (5-HT<sub>2c</sub>R agonist) | Time on top (s) | Treatment [1, 114] | 6.337 | 0.0132 | 0.039 |
|  |  |  | Dose [2, 114] | 3.665 | 0.0287 | 0.039 |
|  |  |  | Interaction [2, 114] | 4.602 | 0.012 | 0.052 |
| | | Time on bottom (s) | Treatment [1,114] | 20.995 | $1.18 \times 10^{-5}$ | 0.12 |
| | | | Dose [2, 114] | 11.455 | $2.93 \times 10^{-5}$ | 0.125 |
|  |  |  | Interaction [2, 114] | 4.085 | 0.0194 | 0.037 |
| | | Erratic swimming (abs. turn angle) | Treatment [1,114] | 5.137 | 0.0253 | $\omega^2 = 0.017$ |
|  |  |  | Dose [2, 114] | 2.948 | 0.0565 | 0.016 |
| | | | Interaction [2, 114] | 57.101 | $< 2 \times 10^{-16}$ | 0.467 |
|  |  | Freezing (s) | Treatment [1,114] | 15.219 | 0.000162 | 0.105 |
|  |  |  | Dose [2, 114] | 1.037 | 0.358 | 0.001 |
|  |  |  | Interaction [2, 114] | 1.662 | 0.194 | 0.01 |
|  |  | Swimming speed (cm/s) | Treatment [1,114] | 0.005 | 0.9459 | -0.008 |
|  |  |  | Dose [2, 114] | 0.211 | 0.8105 | 0.013 |
|  |  |  | Interaction [2, 114] | 4.652 | 0.0114 | 0.059 |

Post-hoc tests revealed that CAS increased time on bottom (p < 0.001, d = −0.77 for the main effect), and the highest MK-212 dose blocked this effect (*1 mg/kg:* p = 0.39, d = −0.64, non-treated controls vs. treated CAS; p = 0.38, d = 0.6, non-treated CAS vs. treated CAS; *2 mg/kg:* p = 0.828, d = 0.4, non-treated controls vs. treated CAS; p < 0.001, d = 1.64, non-treated CAS vs. treated CAS; Figure 2B).

Post-hoc tests showed that CAS greatly increased erratic swimming (absolute turn angle) in non-treated animals (p < 0.001, d = −2.58, non-treated controls vs. non-treated CAS), an effect that was blocked by both MK-212 doses (*1 mg/kg*: p = 1, d = −0.04, non-treated controls vs. treated CAS; p < 0.001, d = 2.53, non-treated CAS vs. treated CAS; *2 mg/kg*: p = 0.81, d = −0.42, non-treated controls vs. treated CAS; p < 0.001, d = 2.16, non-treated CAS vs. treated CAS; Figure 2C); however, MK-212 also increased absolute turn angle in animals which were not exposed to CAS (*1 mg/kg*: p < 0.001, d = −1.95, non-treated controls vs. treated controls; *2 mg/kg:* p < 0.001, d = −1.91, non-treated controls vs. treated controls).

Post-hoc tests suggested that CAS increased freezing at all MK-212 doses and controls (p < 0.001, d = −0.72 for the main effect of CAS; Figure 2D).

While a small-to-medium-sized interaction effect was found for swimming speed (Table 1), post-hoc tests found no differences across groups (Figure 2E).

#### 3.1.2. After exposure

Table 2 presents full ANOVA results for effects in the novel tank test after CAS exposure and washout. Post-hoc tests revealed that MK-212 (1 mg/kg) increased time on top in both controls and CAS-exposed animals (*1 mg/kg*: p = 0.019, d = −0.63 vs. non-treated animals; p = 0.038, d = 0.54 vs. 2 mg/kg; *2 mg/kg:* p = 0.92, d = −0.09 vs. non-treated animals; Figure 3A).

**Figure 3.**
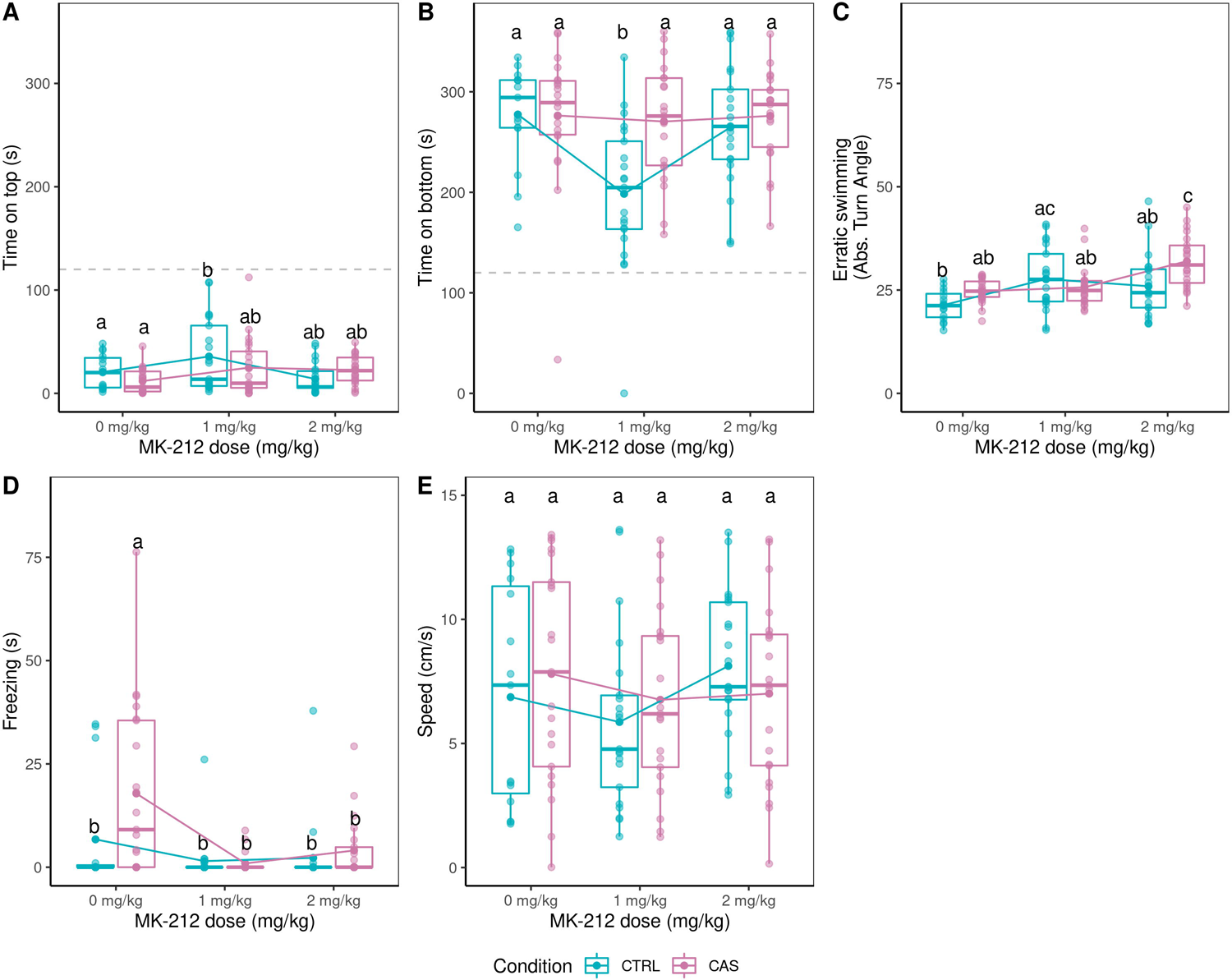
Effects of MK-212 on behavior <u>after</u> CAS exposure. (A) Time spent on top third of the tank. (B) Time spent on bottom third of the tank. (C) Erratic swimming. (D) Total time spent freezing. (E) Swimming speed. Different letters represent statistical differences at the p < 0.05 level; similar letters indicate lack of statistically significant differences. Data are presented as individual data points (dots) superimposed over the median ± interquartile ranges. Dashed lines on panels A and B represent change levels. Dots connected by lines represent group means. CTRL = controls (water-exposed animals); CAS = conspecific alarm substance. Final sample sizes: CTRL + VEH: n = 15 animals; CTRL + 1 mg/kg MK-212: n = 21 animals; CTRL + 2 mg/kg MK-212: n = 21 animals; CAS + VEH: n = 21 animals; CAS + 1 mg/kg MK-212: n = 21 animals; CAS + 2 mg/kg MK-212: n = 21 animals.

**Table 2.** Full ANOVA results for effects of MK-212 on behavioral endpoints after CAS exposure.

| Context | Drug | Endpoint | Effect<br>[df, df2] | F | p-value | Effect size<br>($\omega^2$) |
| --- | --- | --- | --- | --- | --- | --- |
| After exposure | MK-212 (5-HT<sub>2c</sub>R agonist) | Time on top (s) | Treatment [1, 114] | 0.945 | 0.333 | 0.0 |
|  |  |  | Dose [2, 114] | 4.892 | 0.009 | 0.06 |
|  |  |  | Interaction [2, 114] | 2.393 | 0.0959 | 0.021 |
|  |  | Time on bottom (s) | Treatment [1,114] | 7.751 | 0.0063 | 0.048 |
|  |  |  | Dose [2, 114] | 5.385 | 0.0058 |  |
|  |  |  | Interaction [2, 114] | 4.203 | 0.0173 | 0.045 |
|  |  | Erratic swimming (abs. turn angle) | Treatment [1,114] | 3.641 | 0.0589 | 0.018 |
|  |  |  | Dose [2, 114] | 9.244 | 0.00019 | 0.112 |
|  |  |  | Interaction [2, 114] | 4.951 | 0.00867 | 0.054 |
|  |  | Freezing (s) | Treatment [1,114] | 4.658 | 0.033 | 0.025 |
|  |  |  | Dose [2, 114] | 11.617 | 2.56 x 10<sup>-5</sup> | 0.143 |
|  |  |  | Interaction [2, 114] | 2.777 | 0.0664 | 0.024 |
|  |  | Swimming speed (cm/s) | Treatment [1,114] | 0.111 | 0.74 | -0.007 |
|  |  |  | Dose [2, 114] | 1.339 | 0.266 | 0.006 |
|  |  |  | Interaction [2, 114] | 0.981 | 0.378 | 0.0 |

Post-hoc tests suggested that MK-212 (1 mg/kg) decreased time on bottom in control animals (Figure 3B), but not in CAS-exposed animals (*1 mg/kg*: p = 0.002, d = 1.31, non-treated controls vs. 1 mg/kg controls; p = 1.0, d = 0.1, non-treated CAS vs. 1 mg/kg CAS; *2 mg/kg:* p = 0.99, d = 0.21, non-treated controls vs. 2 mg/kg controls; p = 1, d = 0.01, non-treated CAS vs. 2 mg/kg CAS).

Post-hoc tests suggested a synergistic effect between CAS and MK-212 at 2 mg/kg, which potentiated CAS-elicited increases in erratic swimming (Figure 3C); moreover, 1 mg/kg MK-212 increased erratic swimming in control animals (*1 mg/kg:* p = 0.026, d= −1.05, non-treated controls vs. 1 mg/kg controls; p = 1.0, d = −0.16, non-treated CAS vs. 1 mg/kg CAS; *2 mg/kg:* p = 0.22, d = −0.76, non-treated controls vs. 2 mg/kg controls; p = 0.002, d = −1.21, non-treated CAS vs. 2 mg/kg CAS).

Post-hoc tests revealed that CAS increased freezing (p = 0.05, d = −0.364 for the main effect), an effect that was blocked by all MK-212 doses (*1 mg/kg:* p = 0.74, d = 0.47, non-treated controls vs. 1 mg/kg controls; p < 0.001, d = 1.5, non-treated CAS vs. 1 mg/kg CAS; *2 mg/kg:* p = 0.85, d = 0.4, non-treated controls vs. 2 mg/kg controls; p = 0.002, d = 1.22, non-treated CAS vs. 2 mg/kg CAS; Figure 3D).

No effects were found for swimming speed (Figure 3E).

### 3.2. Effects of WAY-161503 on alarm reaction and post-exposure behavior

#### 3.2.1. During exposure

No outliers were detected from any group in this experiment. Table 3 presents full ANOVA results for this experiment. Post-hoc tests showed that while CAS did not alter time on top (p = 0.054, d = 1.0, non-treated controls vs. non-treated CAS), a synergistic effect was apparent, with WAY-161503-treated animals showing less time on top than controls (p = 0.381, d = 0.618, non-treated controls vs. WAY-161503 controls; p = 0.977, d = 0.125, non-treated CAS vs. WAY-161503 CAS; p = 0.023, d = 1.125, non-treated controls vs. WAY 161-503 CAS; Figure 4A).

**Figure 4.**
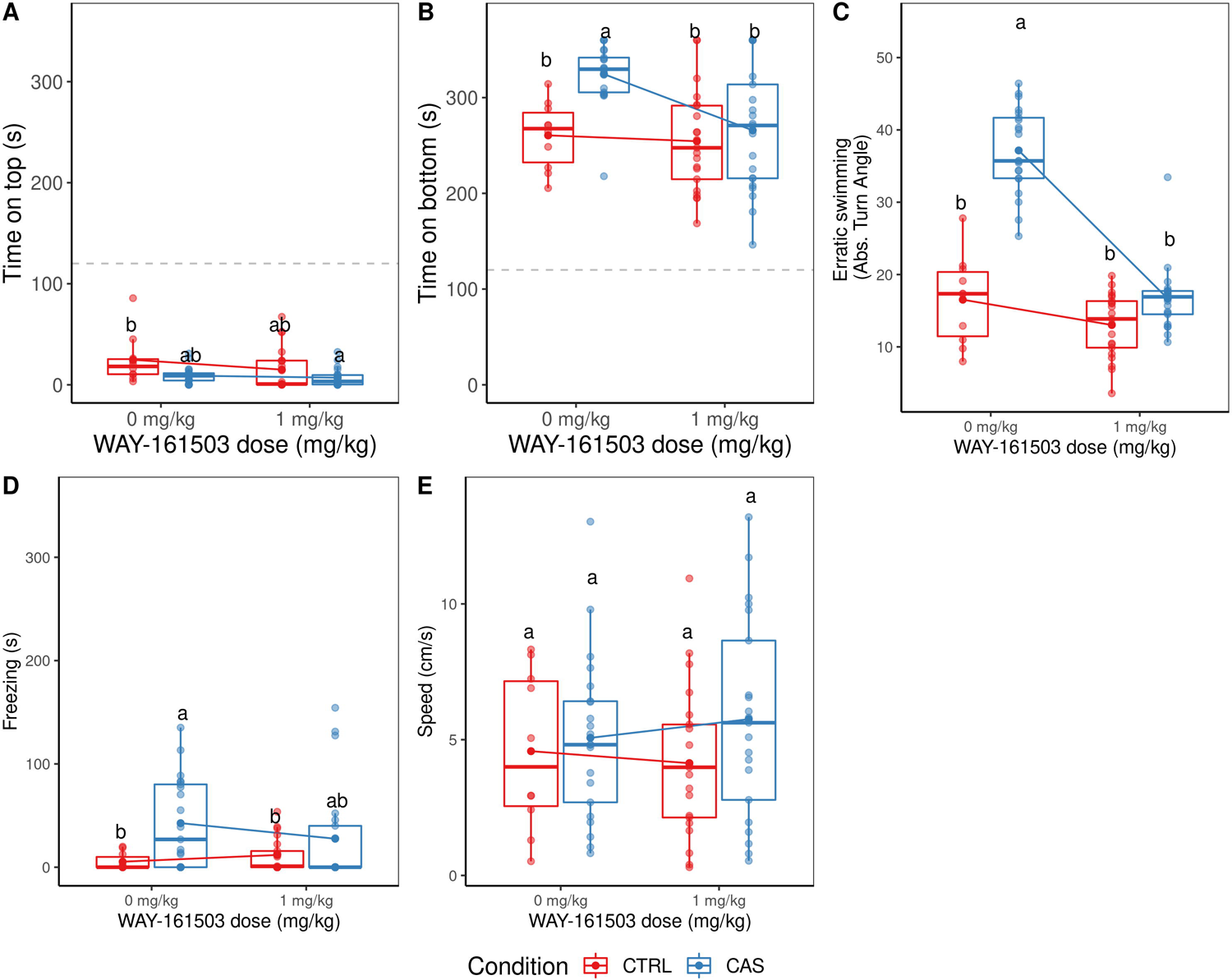
Effects of WAY-161503 on behavior <u>during</u> CAS exposure. (A) Time spent on top third of the tank. (B) Time spent on bottom third of the tank. (C) Erratic swimming. (D) Total time spent freezing. (E) Swimming speed. Different letters represent statistical differences at the p < 0.05 level; similar letters indicate lack of statistically significant differences. Data are presented as individual data points (dots) superimposed over the median ± interquartile ranges. Dashed lines on panels A and B represent change levels. Dots connected by lines represent group means. CTRL = controls (water-exposed animals); CAS = conspecific alarm substance. Final sample sizes: CTRL + VEH: n = 10 animals; CTRL + 1 mg/kg WAY-161503: n = 21 animals; CAS + VEH: n = 21 animals; CAS + 1 mg/kg WAY-161503: n = 21 animals.

**Table 3.** Full ANOVA results for effects of WAY-161503 on behavioral endpoints during CAS exposure.

| Context | Drug | Endpoint | Effect<br>[df1, df2] | F | P-value | Effect size<br>($\omega^2$) |
| --- | --- | --- | --- | --- | --- | --- |
| During exposure | WAY-161503 (5-HT<sub>2c</sub>R agonist) | Time on top (s) | Treatment [1, 69] | 7.404 | 0.00823 | 0.08 |
|  |  |  | Drug [1, 69] | 1.751 | 0.19011 | 0.009 |
|  |  |  | Interaction [1, 69] | 1.001 | 0.32067 | 0.0 |
|  |  | Time on bottom (s) | Treatment [1, 69] | 10.842 | 0.00157 | 0.103 |
|  |  |  | Drug [1, 69] | 10.138 | 0.00218 | 0.096 |
|  |  |  | Interaction [1, 69] | 4.614 | 0.03523 | 0.038 |
|  |  | Erratic swimming (abs. turn angle) | Treatment [1, 69] | 108.37 | < 0.0001 | 0.311 |
|  |  |  | Drug [1, 69] | 122.99 | < 0.0001 | 0.353 |
|  |  |  | Interaction [1, 69] | 44.24 | < 0.0001 | 0.125 |
|  |  | Freezing (s) | Treatment [1, 69] | 8.582 | 0.0046 | 0.094 |
|  |  |  | Drug [1, 69] | 0.564 | 0.4552 | -0.005 |
|  |  |  | Interaction [1, 69] | 1.434 | 0.2352 | 0.005 |
|  |  | Swimming speed (cm/s) | Treatment [1, 69] | 2.303 | 0.134 | 0.018 |
|  |  |  | Drug [1, 69] | 0.103 | 0.749 | -0.012 |
|  |  |  | Interaction [1, 69] | 0.527 | 0.471 | -0.006 |

Post-hoc tests revealed that CAS increased time on bottom (p = 0.007, d = −1.287, non-treated controls vs. non-treated CAS), and WAY-161503 blocked this effect (p = 0.989, d = 0.123, non-treated controls vs. WAY-161503 controls; p = 0.002, d = 1.181, non-treated CAS vs. WAY-161503 CAS; Figure 4B).

Again, post-hoc tests suggested that CAS increased erratic swimming (p < 0.001, d = − 3.984, non-treated controls vs. non-treated CAS), while WAY-161503 blocked this effect (p = 0.302, d = 0.6759, non-treated controls vs. WAY-161503 controls; p < 0.001, d = 3.9537, non-treated CAS vs. WAY-161503 CAS; Figure 4C).

Post-hoc tests revealed that CAS increased freezing (p = 0.047, d = −1.02, non-treated controls vs. non-treated CAS), but this effect was not blocked by WAY-161503 (p = 0.967, d = − 0.178, non-treated controls vs. WAY-161503 controls; p = 0.544, d = 0.412, non-treated CAS vs. WAY-161503 CAS; Figure 4D).

Finally, no effects were found for swimming speed (Figure 4E).

#### 3.2.2. After exposure

Full ANOVA results for this experiment can be found on Table 4. Post-hoc effects revealed a synergistic effect for time on top, with WAY-161503 increasing time on top in animals exposed to CAS (p = 0.922, d = 0.242, non-treated controls vs. non-treated CAS; p = 0.907, 0.259, non-treated controls vs. WAY-161503 controls; p = 0.011, d = 0.9833, non-treated CAS vs. WAY-161503 CAS; Figure 5A).

**Figure 5.**
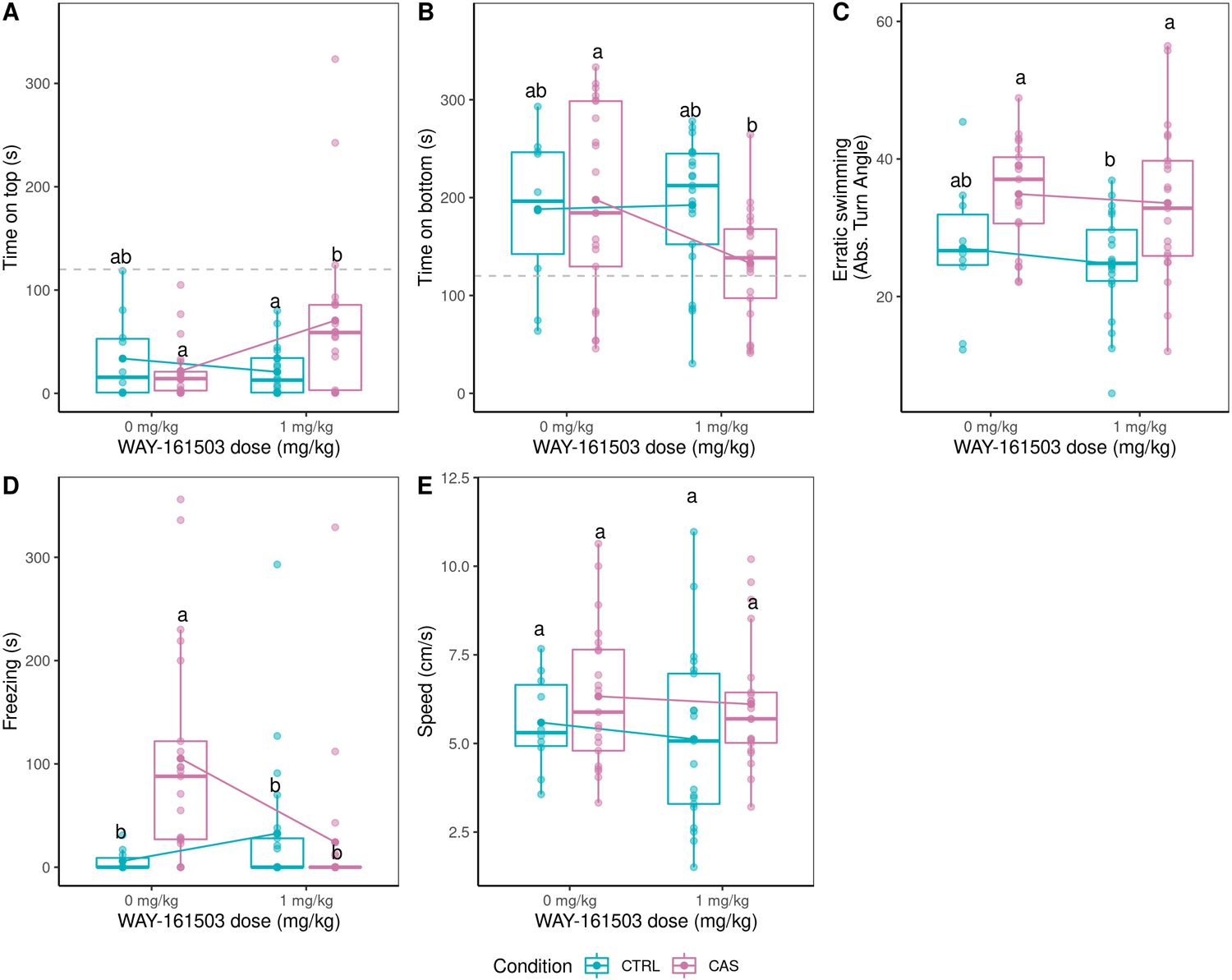
Effects of WAY-161503 on behavior <u>after</u> CAS exposure. (A) Time spent on top third of the tank. (B) Time spent on bottom third of the tank. (C) Erratic swimming. (D) Total time spent freezing. (E) Swimming speed. Different letters represent statistical differences at the p < 0.05 level; similar letters indicate lack of statistically significant differences. Data are presented as individual data points (dots) superimposed over the median ± interquartile ranges. Dashed lines on panels A and B represent change levels. Dots connected by lines represent group means. CTRL = controls (water-exposed animals); CAS = conspecific alarm substance. Final sample sizes: CTRL + VEH: n = 10 animals; CTRL + 1 mg/kg WAY-161503: n = 21 animals; CAS + VEH: n = 21 animals; CAS + 1 mg/kg WAY-161503: n = 21 animals.

**Table 4.** Full ANOVA results for effects of WAY-161503 on behavioral endpoints after CAS exposure.

| Context | Drug | Endpoint | Effect<br>[df1, df2] | F | P-value | Effect size<br>($\omega^2$) |
| --- | --- | --- | --- | --- | --- | --- |
| After exposure | WAY-161503 (5-HT<sub>2C</sub>R agonist) | Time on top (s) | Treatment [1, 69] | 3.221 | 0.0771 | 0.026 |
|  |  |  | Drug [1, 69] | 4.254 | 0.0429 | 0.039 |
|  |  |  | Interaction [1, 69] | 0.039<br>6.352 | 0.014 | 0.064 |
|  |  | Time on bottom (s) | Treatment [1, 69] | 1.95 | 0.1671 | 0.012 |
|  |  |  | Drug [1, 69] | 4.107 | 0.0466 | 0.039 |
|  |  |  | Interaction [1, 69] | 3.267 | 0.0751 | 0.029 |
|  |  | Erratic swimming (abs. turn angle) | Treatment [1, 69] | 16.119 | 0.000149 | 0.174 |
|  |  |  | Drug [1, 69] | 0.568 | 0.454 | -0.005 |
|  |  |  | Interaction [1, 69] | 0.042 | 0.838 | -0.011 |
|  |  | Freezing (s) | Treatment [1, 69] | 4.726 | 0.0331 | 0.43 |
|  |  |  | Drug [1, 69] | 4.17 | 0.04497 | 0.037 |
|  |  |  | Interaction [1, 69] | 7.663 | 0.00723 | 0.077 |
|  |  | Swimming speed (cm/s) | Treatment [1, 69] | 3.85 | 0.0538 | 0.038 |
|  |  |  | Drug [1, 69] | 0.416 | 0.5212 | -0.008 |
|  |  |  | Interaction [1, 69] | 0.062 | 0.8037 | -0.013 |

Post-hoc tests suggested a synergistic effect also on time on bottom, with WAY-161503 decreasing time on bottom in animals exposed to CAS (p = 0.988, d = −0.1243, non-treated controls vs. non-treated CAS; p = 0.999, d = −0.0538, non-treated controls vs. WAY-161503 controls; p = 0.041, d = 0.8369, non-treated CAS vs. WAY-161503 CAS; Figure 5B).

While ANOVA found effects of treatment on erratic swimming, post-hoc tests did not find an increase in erratic swimming with CAS (p = 0.125, d = −0.857), nor a synergistic effect of WAY-161503 (p = 0.922, d = 0.243, non-treated controls vs. WAY-161503 controls; p = 0.968, d = 0.142, non-treated CAS vs. WAY-161503 CAS; Figure 5C).

Post-hoc tests revealed that CAS increased freezing (p = 0.009, d = −1.257, non-treated controls vs. non-treated CAS), while WAY-161503 blocked this effect (p = 0.816, d = −0.338, non-treated controls vs. WAY-161503 controls; p = 0.008, d = 1.026, non-treated CAS vs. WAY-161-503 CAS; Figure 5D).

Again, no effects were found for swimming speed (Figure 5E).

### 3.3. Effects of RS-102221 on alarm reaction and post-exposure behavior

#### 3.3.1. During exposure

No outliers were detected in any groups. Full results for ANOVAs can be found on Table 5. Post-hoc tests suggested that CAS decreased time on top (p = 0.004, d = 0.74 for the main effect); RS-102221 was not able to change this effect (p = 0.353, d = 0.64, non-treated controls vs. treated CAS; p = 0.941, d = −0.18, non-treated CAS vs. treated CAS; Figure 6A).

**Figure 6.**
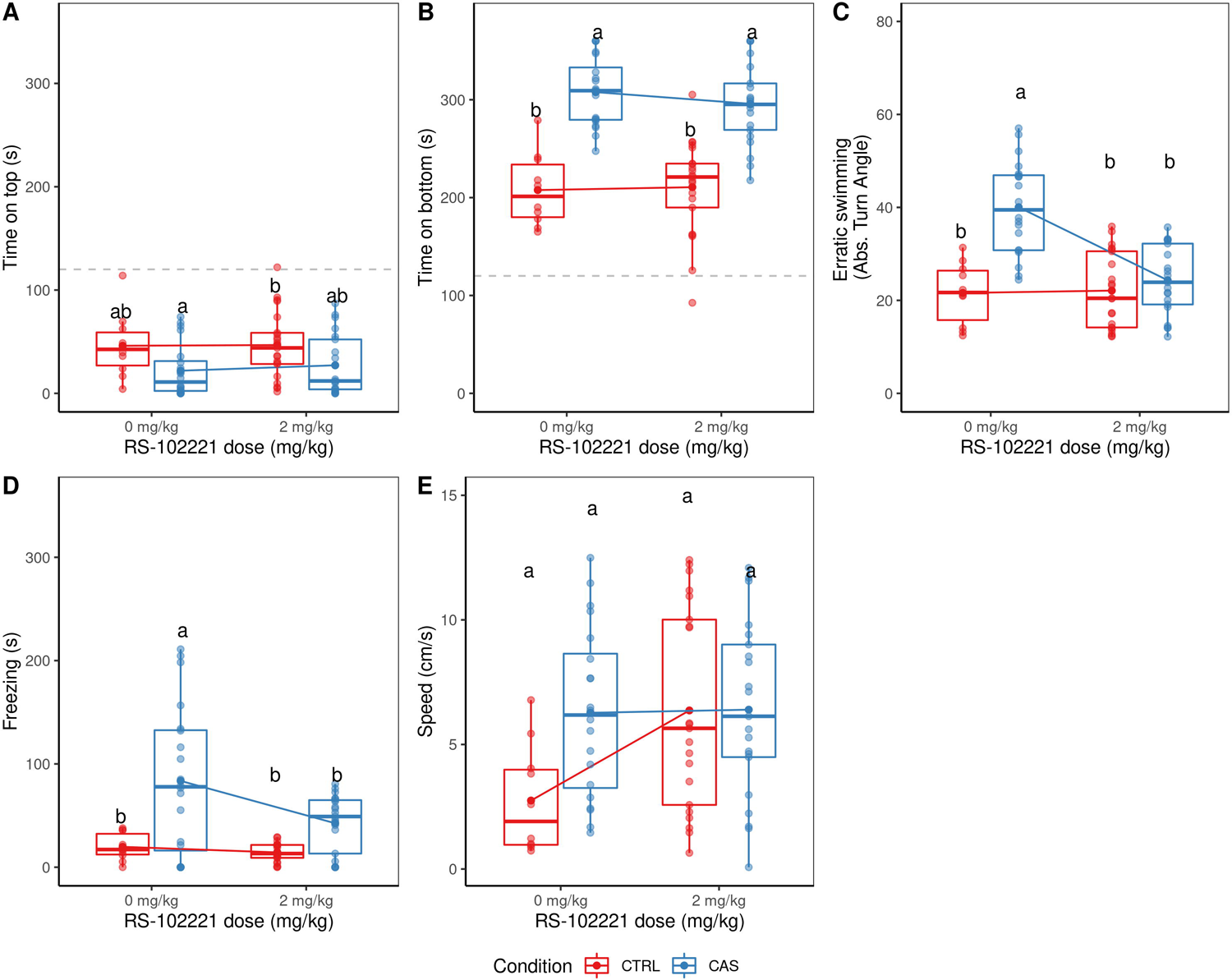
Effects of RS-102221 on behavior <u>during</u> CAS exposure. (A) Time spent on top third of the tank. (B) Time spent on bottom third of the tank. (C) Erratic swimming. (D) Total time spent freezing. (E) Swimming speed. Different letters represent statistical differences at the p < 0.05 level; similar letters indicate lack of statistically significant differences. Data are presented as individual data points (dots) superimposed over the median ± interquartile ranges. Dashed lines on panels A and B represent change levels. Dots connected by lines represent group means. CTRL = controls (water-exposed animals); CAS = conspecific alarm substance. Final sample sizes: CTRL + VEH: n = 10 animals; CTRL + 2 mg/kg RS-102221: n = 21 animals; CAS + VEH: n = 20 animals; CAS + 2 mg/kg RS-102221: n = 21 animals.

**Table 5.** Full ANOVA results for effects of RS-102221 on behavioral endpoints during CAS exposure.

| Context | Drug | Endpoint | Effect<br>[df1, df2] | F | P-value | Effect size<br>($\omega^2$) |
| --- | --- | --- | --- | --- | --- | --- |
| During exposure | RS-102221 (5-HT<sub>2C</sub>R antagonist) | Time on top (s) | Treatment [1, 68] | 9.68 | 0.003 | 0.011 |
|  |  |  | Drug [1, 68] | 0.231 | 0.632 | -0.01 |
|  |  |  | Interaction [1, 68] | 0.096 | 0.757 | -0.012 |
|  |  | Time on bottom (s) | Treatment [1, 68] | 85.978 | < 0.0001 | 0.53 |
|  |  |  | Drug [1, 68] | 0.352 | 0.555 | -0.009 |
|  |  |  | Interaction [1, 68] | 0.517 | 0.474 | -0.006 |
|  |  | Erratic swimming (abs. turn angle) | Treatment [1, 68] | 25.74 | < 0.0001 | 0.26 |
|  |  |  | Drug [1, 68] | 21.06 | < 0.0001 | 0.22 |
|  |  |  | Interaction [1, 68] | 15.47 | < 0.0001 | 0.17 |
|  |  | Freezing (s) | Treatment [1, 68] | 21.973 | < 0.0001 | 0.23 |
|  |  |  | Drug [1, 68] | 7.06 | 0.0098 | 0.08 |
|  |  |  | Interaction [1, 68] | 2.98 | 0.089 | 0.03 |
|  |  | Swimming speed (cm/s) | Treatment [1, 68] | 1.84 | 0.179 | 0.01 |
|  |  |  | Drug [1, 68] | 3.156 | 0.08 | 0.03 |
|  |  |  | Interaction [1, 68] | 2.981 | 0.089 | 0.03 |

Post-hoc tests suggested that CAS increased time on bottom (p < 0.001, d = −2.22 for the main effect), and RS-102221 was not able to change this effect (p < 0.001, d = −2.12, non-treated controls vs. treated CAS; p = 0.8, d = 0.29, non-treated CAS vs. treated CAS).

Post-hoc tests suggested that CAS increased erratic swimming (p < 0.01, d = 1.24for the main effect), an effect that was blocked by RS-102221 (p =0.829, d = −0.33, non-treated controls vs. treated CAS; p <0.001, d = 1.89, non-treated CAS vs. treated CAS; Figure 6C).

Post-hoc tests revealed that CAS increased freezing (p < 0.001, d = 1.10 for the main effect), and RS-102221 partially blocked this effect (p = 0.488, d = −0.548, non-treated controls vs. treated CAS; p = 0.013, d = 0.985, non-treated CAS vs. treated CAS; Figure 6D).

Finally, no effects were found for swimming speed (Figure 6E).

#### 3.3.2. After exposure

Full results for ANOVAs for this experiment can be found on Table 6. No effects were found for time on top (Figure 7A); Post-hoc tests suggested that CAS increased time on bottom after exposure (p = 0.05, d = 1.02, non-treated controls vs. non-treated CAS), an effect that was blocked by RS-102221 (p = 0.506, d = 0.54, non-treated controls vs. treated CAS; p < 0.001, d = 1.55, non-treated CAS vs. treated CAS)(Figure 7B).

**Figure 7.**
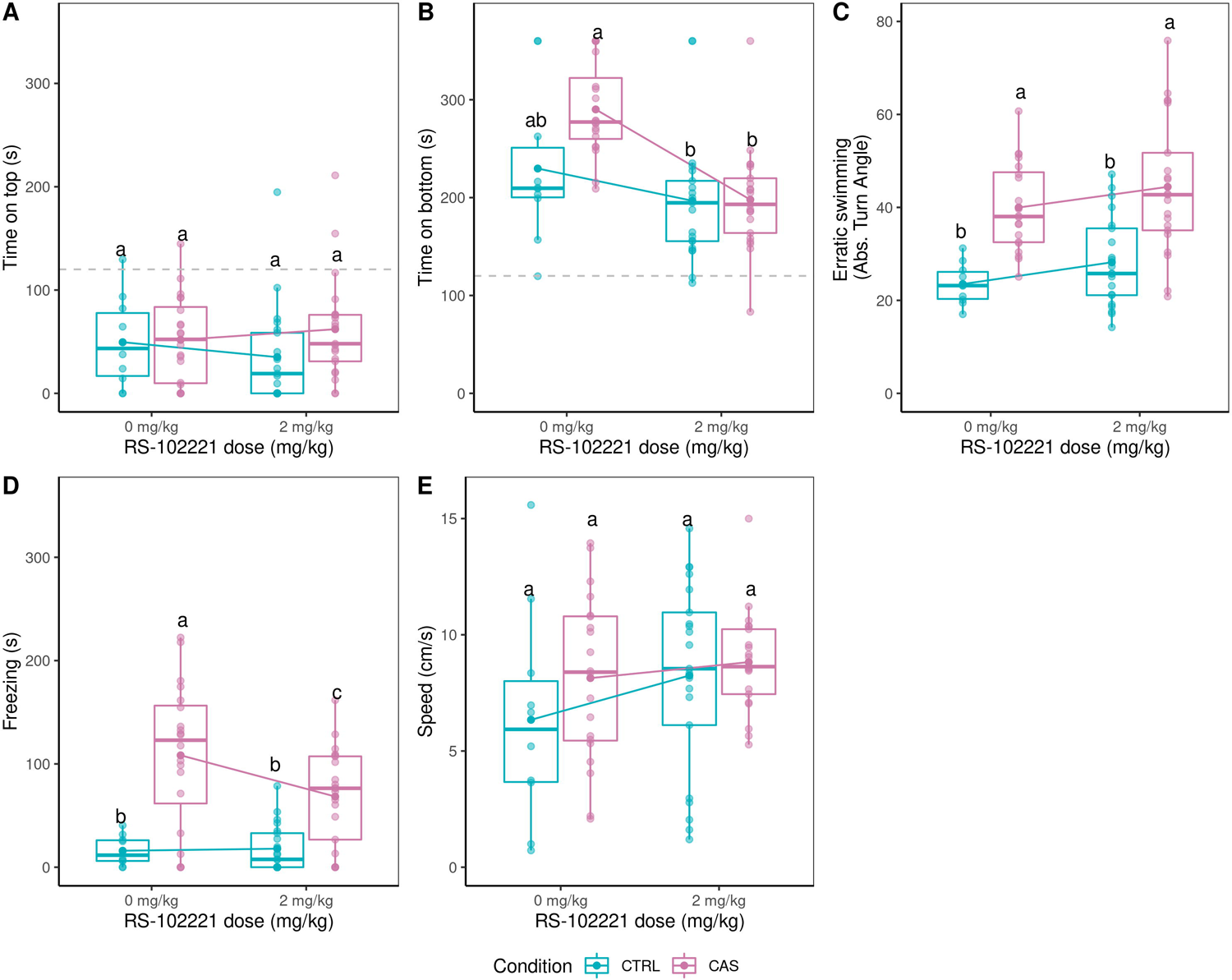
Effects of RS-102221 on behavior <u>after</u> CAS exposure. (A) Time spent on top third of the tank. (B) Time spent on bottom third of the tank. (C) Erratic swimming. (D) Total time spent freezing. (E) Swimming speed. Different letters represent statistical differences at the p < 0.05 level; similar letters indicate lack of statistically significant differences. Data are presented as individual data points (dots) superimposed over the median ± interquartile ranges. Dashed lines on panels A and B represent change levels. Dots connected by lines represent group means. CTRL = controls (water-exposed animals); CAS = conspecific alarm substance. Final sample sizes: CTRL + VEH: n = 10 animals; CTRL + 2 mg/kg RS-102221: n = 21 animals; CAS + VEH: n = 20 animals; CAS + 2 mg/kg RS-102221: n = 21 animals.

**Table 6.** Full ANOVA results for effects of RS-102221 on behavioral endpoints after CAS exposure.

| Context | Drug | Endpoint | Effect<br>[df1, df2] | F | P-value | Effect size<br>($\omega^2$) |
| --- | --- | --- | --- | --- | --- | --- |
| After exposure | RS-102221 (5-HT<sub>2C</sub>R antagonist) | Time on top (s) | Treatment [1, 68] | 2.449 | 0.122 | 0.02 |
|  |  |  | Drug [1, 68] | 0.003 | 0.969 | -0.01 |
|  |  |  | Interaction [1, 68] | 1.152 | 0.287 | 0.002 |
|  |  | Time on bottom (s) | Treatment [1, 68] | 6.322 | 0.0143 | 0.07 |
|  |  |  | Drug [1, 68] | 22.796 | < 0.0001 | 0.23 |
|  |  |  | Interaction [1, 68] | 4.08 | 0.047 | 0.04 |
|  |  | Erratic swimming (abs. turn angle) | Treatment [1, 68] | 36.19 | < 0.0001 | 0.33 |
|  |  |  | Drug [1, 68] | 2.957 | 0.09 | 0.03 |
|  |  |  | Interaction [1, 68] | 0.002 | 0.965 | -0.01 |
|  |  | Freezing (s) | Treatment [1, 68] | 39.763 | < 0.0001 | 0.35 |
|  |  |  | Drug [1, 68] | 4.16 | 0.045 | 0.04 |
|  |  |  | Interaction [1, 68] | 3.23 | 0.077 | 0.03 |
|  |  | Swimming speed (cm/s) | Treatment [1, 68] | 1.014 | 0.317 | 0.0002 |
|  |  |  | Drug [1, 68] | 1.83 | 0.181 | 0.01 |
|  |  |  | Interaction [1, 68] | 0.475 | 0.493 | -0.008 |

Post-hoc tests suggested that CAS increased erratic swimming after exposure (p = 0.001, d = 1.514, non-treated controls vs. non-treated CAS), an effect that was not altered by RS-102221 (p < 0.001, d = −1.92, non-treated controls vs. treated CAS; p = 0.56, d = −0.41, non-treated CAS vs. treated CAS)(Figure 7C).

Post-hoc tests suggested that CAS increased freezing after exposure (p < 0.001, d = 1.96, non-treated controls vs. non-treated CAS), an effect that was partially blocked by RS-102221 (p = 0.025, d = −1.12, non-treated controls vs. treated CAS; p = 0.041, d = 0.85; Figure 7D).

Finally, no effects were found for swimming speed (Figure 7E).

### 3.4. Restraint stress-elicited behavioral changes

#### 3.4.1. Effects of MK-212 on restraint stress-elicited behavioral changes

Full results for ANOVAs for this experiment can be found on Table 7. No effects were found for time on top (Figure 8A). Post-hoc tests showed that ARS increased time on bottom (*p* = 0.038, *d* = −1.566 vs. non-treated controls), and MK-212 (2 mg/kg) blocked this effect (*p* = 0.757, *d* = −0.495 non-treated controls vs. treated ARS; Figure 8B Post-hoc tests suggested that a ARS increased erratic swimming (p < 0.001, d = −1.57 for the main effect), an effect that was not blocked by MK-212 (*p* = 0.006, d = −1.85, non-treated controls vs. treated ARS; Figure 8C).

**Figure 8.**
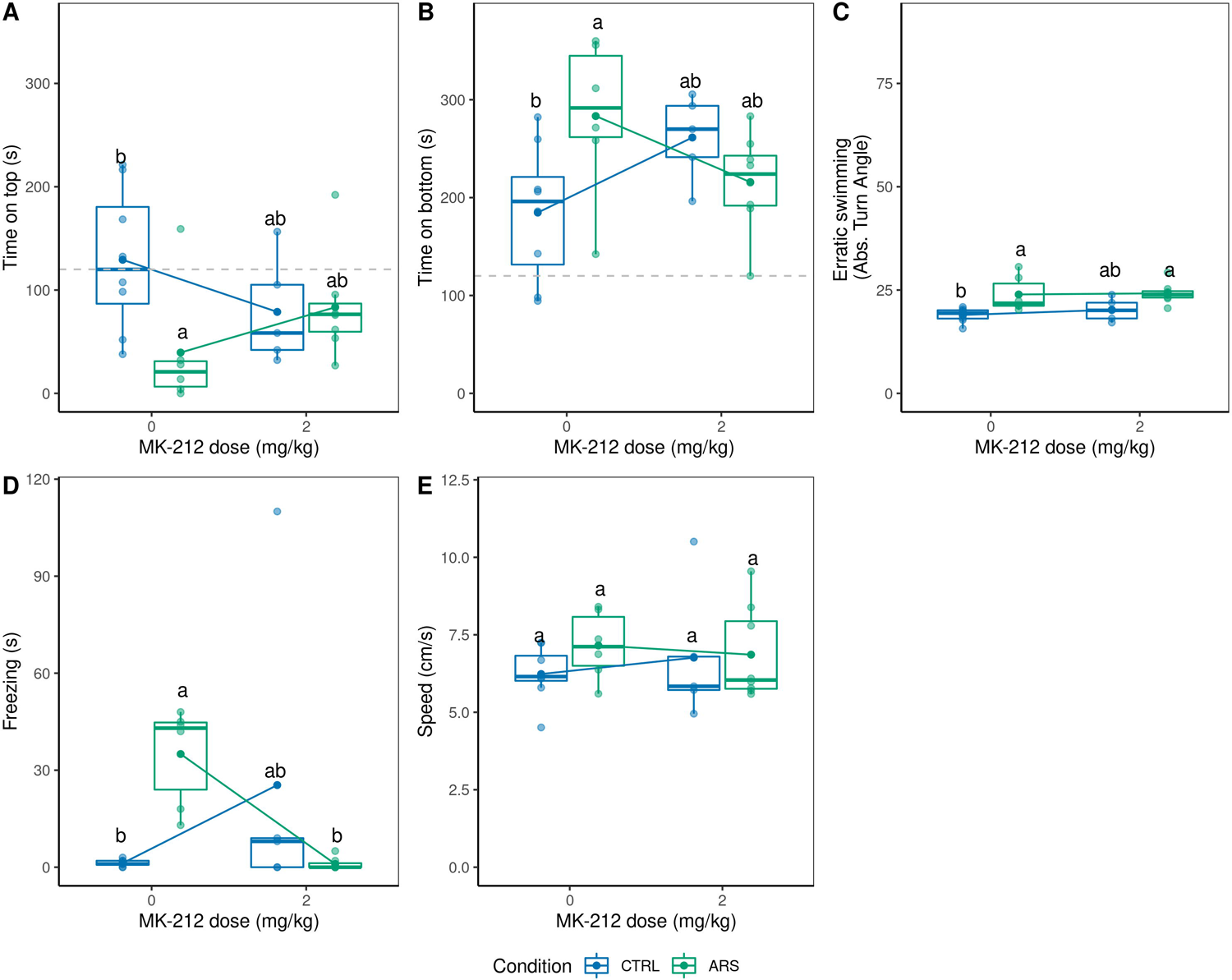
Effects of MK-212 on behavior <u>after</u> acute restraint stress (ARS). (A) Time spent on top third of the tank. (B) Time spent on bottom third of the tank. (C) Erratic swimming. (D) Total time spent freezing. (E) Swimming speed. Different letters represent statistical differences at the p < 0.05 level; similar letters indicate lack of statistically significant differences. Data are presented as individual data points (dots) superimposed over the median ± interquartile ranges. Dashed lines on panels A and B represent change levels. Dots connected by lines represent group means. CTRL = controls (water-exposed animals); ARS = acute restraint stress. Final sample sizes: CTRL + VEH: n = 8 animals; CTRL + 2 mg/kg MK-212: n = 5 animals; ARS + VEH: n = 6 animals; ARS + 2 mg/kg MK-212: n = 8 animals.

**Table 7.** Full ANOVA results for effects of MK-212 on behavioral endpoints after restraint stress.

| Context | Drug | Endpoint | Effect<br>[df1, df2] | F | P-value | Effect size<br>($\omega^2$) |
| --- | --- | --- | --- | --- | --- | --- |
| After restraint stress | MK-212 (5-HT<sub>2c</sub>R agonist) | Time on top (s) | Treatment [1, 23] | 0.167 | 0.687 | -0.026 |
|  |  |  | Drug [1, 23] | 3.884 | 0.061 | 0.089 |
|  |  |  | Interaction [1, 23] | 4.215 | 0.052 | 0.1 |
|  |  | Time on bottom (s) | Treatment [1, 23] | 6.3864 | 0.019 | 0.099 |
|  |  |  | Drug [1, 23] | 0.0879 | 0.77 | -0.017 |
|  |  |  | Interaction [1, 23] | 8.5023 | 0.008 | 0.22 |
|  |  | Erratic swimming (abs. turn angle) | Treatment [1, 23] | 17.9732 | 0.00031 | 0.398 |
|  |  |  | Drug [1, 23] | 0.446 | 0.511 | 0.013 |
|  |  |  | Interaction [1, 23] | 0.193 | 0.664 | -0.019 |
|  |  | Freezing (s) | Treatment [1, 23] | 0.384 | 0.541 | -0.016 |
|  |  |  | Drug [1, 23] | 0.618 | 0.44 | -0.01 |
|  |  |  | Interaction [1, 23] | 12.339 | 0.002 | 0.304 |
|  |  | Swimming speed (cm/s) | Treatment [1, 23] | 1.0005 | 0.328 | 0.0 |
|  |  |  | Drug [1, 23] | 0.0283 | 0.868 | -0.038 |
|  |  |  | Interaction [1, 23] | 0.5437 | 0.468 | -0.018 |

Post-hoc tests suggested that animals injected with MK-212 and subjected to ARS showed decreased freezing in relation to other groups (p = 0.033, d = −1.6, non-treated controls vs. treated ARS; p = 0.031, d = 1.6, non-treated ARS vs. treated ARS; Figure 8D).

Finally, no effects were found for swimming speed (Figure 8E).

#### 3.4.2. Effects of RS-102221 on restraint stress-elicited behavioral changes

Full results for ANOVAs for this experiment can be found in Table 8. ARS increased time on top (p < 0.0001, d = 7.78 for the main effect), and RS-102221 was not able to block this effect (Figure 9A).

**Figure 9.**
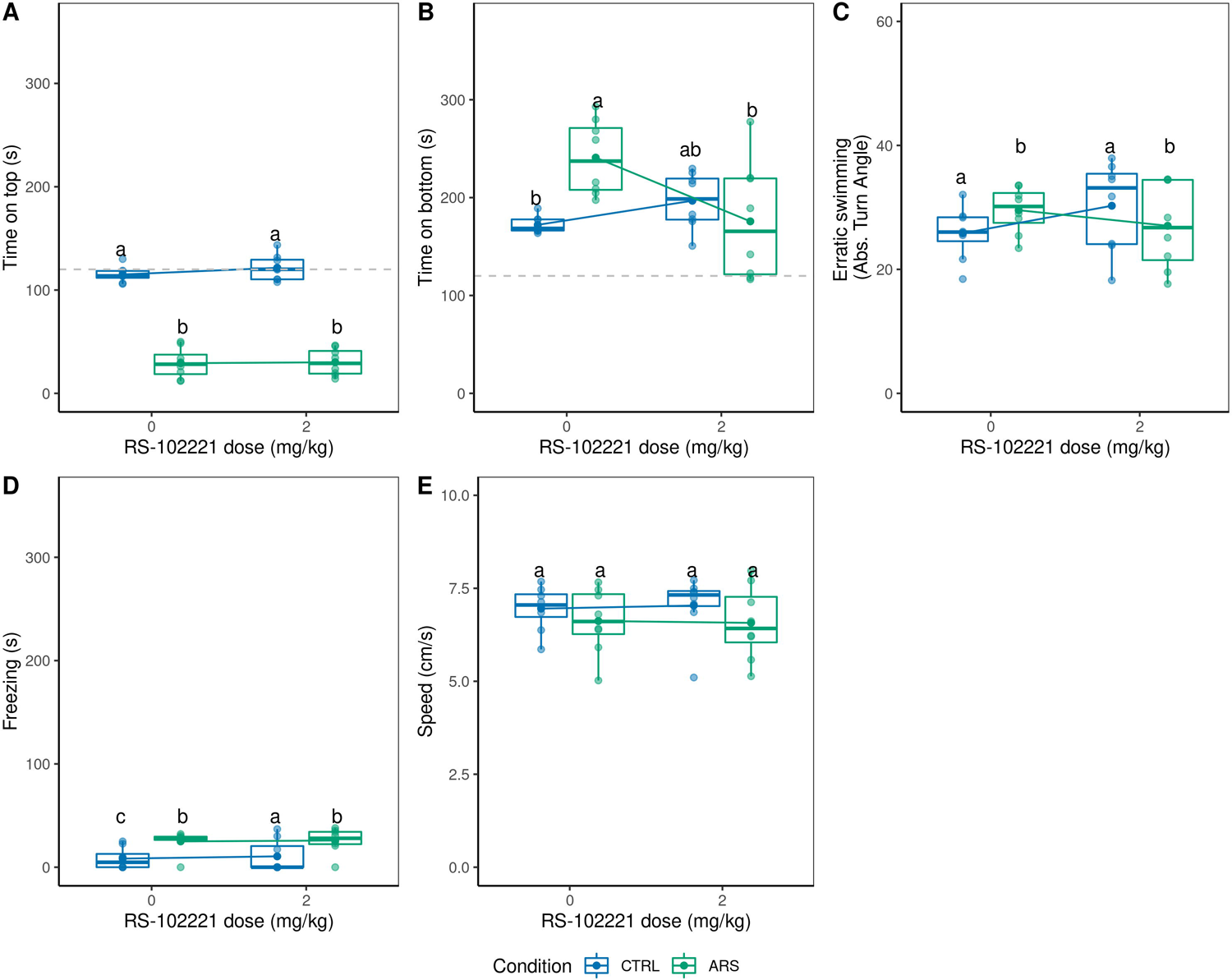
Effects of RS-102221 on behavior <u>after</u> acute restraint stress (ARS). (A) Time spent on top third of the tank. (B) Time spent on bottom third of the tank. (C) Erratic swimming. (D) Total time spent freezing. (E) Swimming speed. Different letters represent statistical differences at the p < 0.05 level; similar letters indicate lack of statistically significant differences. Data are presented as individual data points (dots) superimposed over the median ± interquartile ranges. Dashed lines on panels A and B represent change levels. Dots connected by lines represent group means. CTRL = controls (water-exposed animals); ARS = acute restraint stress. Final sample sizes: CTRL + VEH: n = 8 animals; CTRL + 2 mg/kg RS-102221: n = 8 animals; ARS + VEH: n = 8 animals; ARS + 2 mg/kg RS-102221: n = 8 animals.

**Table 8.**
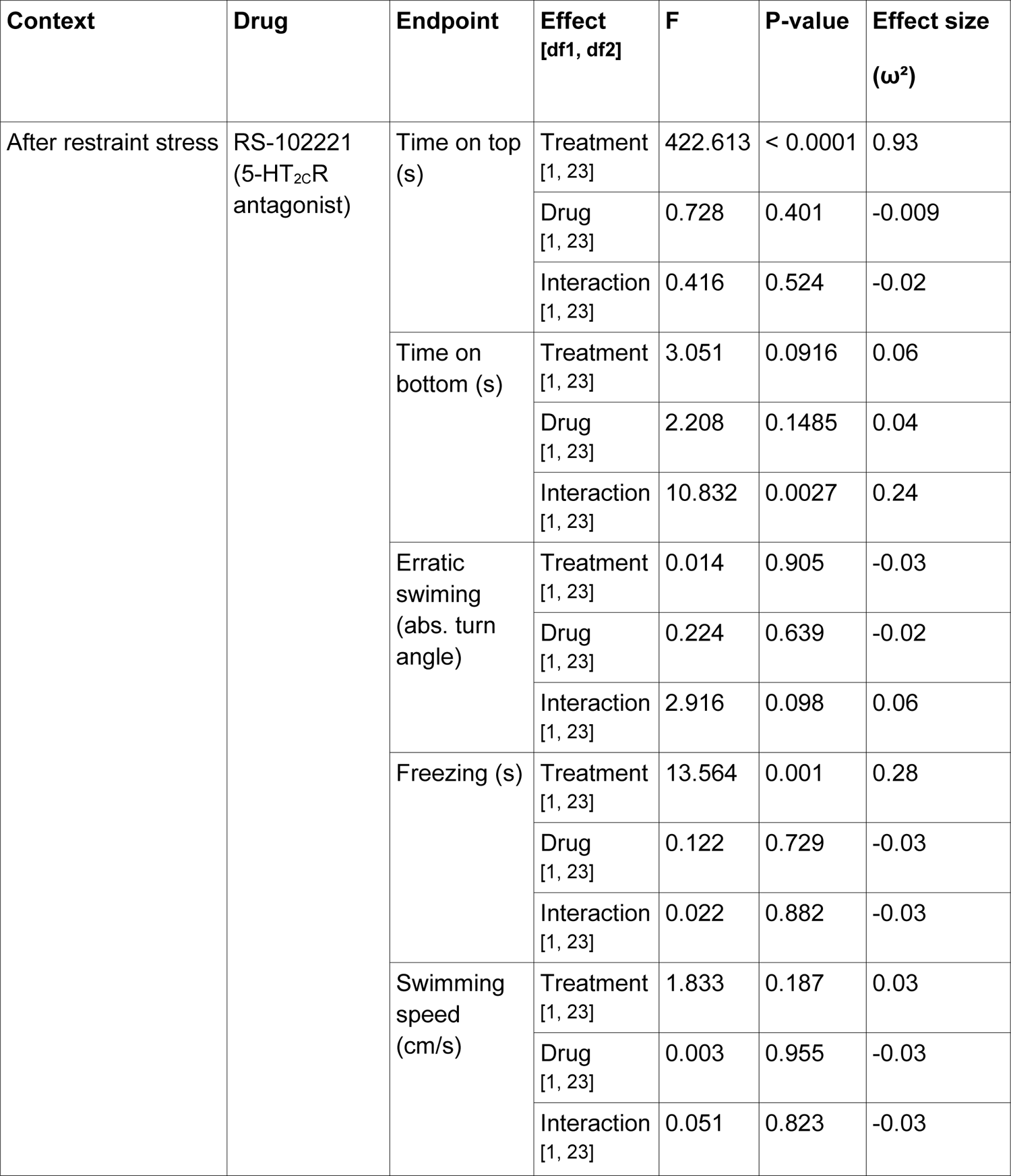
Full ANOVA results for effects of RS-102221 on behavioral endpoints after restraint stress.

Post-hoc tests revealed that ARS increased time on bottom (p = 0.007 vs. non-treated controls), an effect that was partially blocked by RS-102221 (p = 0.997, non-treated controls vs. treated ARS; p = 0.019, non-treated ARS vs. treated ARS; Figure 9B).

No effects were found for erratic swimming (Figure 9C).

ARS increased freezing (p = 0.001, d = 1.4 for the main effect), an effect that was not blocked by RS-102221 (p = 0.052, non-treated controls vs. treated ARS; p = 0.998, non-treated ARS vs. treated ARS; Figure 9D).

Finally, no effects were found for swimming speed (Figure 9E).

### 3.5. Conservation of residues related to ligand binding in zebrafish 5-HT_2C_ receptors

The sequences of *Danio rerio* 5-HT_2C_-like receptors were aligned to the sequence of the human and murine receptors using Clustal Omega. Sequence alignment showing the transmembrane domains is shown in Figure 10. While target residues were conserved across all sequences, differences in residues close to D3.32 the 5-HT_2C_-like receptors zebrafish receptor (S3.30Y in 5-HT_2CL1_ and S3.30F in 5-HT_2CL2_), as well as close to Y7.43 (V7.44I in5-HT_2CL2_) were found.

**Figure 10.**
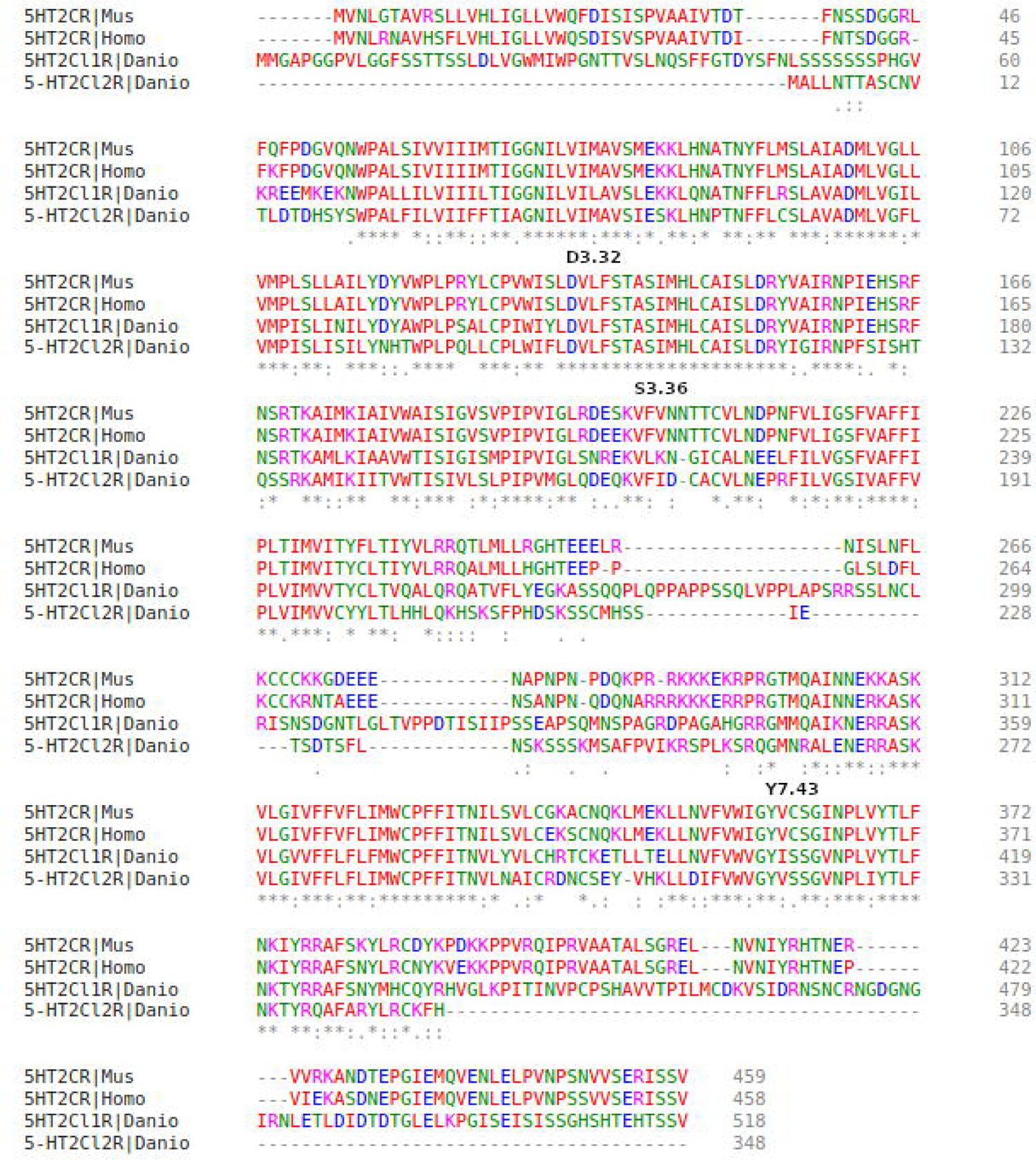
Amino acid sequences for zebrafish 5-HT_2C_RL1 and 5-HT_2C_RL2 aligned with sequences of mice (*Mus*) and humans (*Homo*). Residues associated with the binding site are labeled above the sequences (D3.32, S3.36, Y7.43), according to the Ballesteros et al. (2001) nomenclature. Colors code physicochemical properties: red, small amino acids+hydrophobic amino acids (including aromatic amino acids except tyrosine [Y]); blue, acidic amino acids; magenta, basic amino acids except histidine [H]; green, hydroxyl amino acids+sulfhydryl amino acids+amine+glycine. Asteriks (*) show residues which are conserved across all four sequences, while colons (:) indicate conservation between groups scoring > 0.5 in the Gonnet PAM 250 matrix, and periods (.) indicates conservation between groups scoring < 0.5 in the Gonnet PAM 250 matrix.

## 4. Discussion

The present work tested the hypothesis that phasic activation of the 5-HT_2C_ receptor is involved in behavioral adjustments to distal, but not proximal, threat in zebrafish, and that these receptors exerts a tonic facilitation of defensive behavior to potential threat in zebrafish. We found that 5-HT_2C_ agonists blocked CAS-elicited defensive behavior, but not post-exposure increases in defensive behavior (with the exception of freezing), nor ARS-elicited anxiogenic-like effects. We also found that RS-102221, a 5-HT_2C_ receptor antagonist, did not change behavior during exposure, but it produced a small effect on behavior after exposure to CAS. No effects of RS-102221 were found for ARS-elicited anxiogenic-like effects. Residues related to ligand binding in the human 5-HT_2C_ receptors were conserved in both zebrafish isoforms, but differences exist in residues close to these sites.

### 4.1. Effects of stressors on zebrafish behavior

One of the aims of this paper was to confirm the behavioral effects of CAS during and after exposure (Lima-Maximino et al., 2020), given the considerable variation in the literature (Maximino et al., 2019), and to compare the effects after exposure with those of ARS - considering that, while both CAS and ARS are anxiogenic stressors, from an ecological point of view both should affect different behavioral endpoints.

In the present experiments, CAS consistently increased bottom-dwelling and erratic swimming during exposure, with a smaller component of freezing. After exposure, a strong component of freezing was present, while erratic swimming contributed less to the overall behavioral pattern. These results are consistent with what was observed both with a washout period (Lima-Maximino et al., 2020), and in the absence of a washout, but with an extended observation interval (Mathuru et al., 2012; Nathan et al., 2015). These effects suggest that, as threat levels change from distal (i.e., CAS is present) to potential (i.e., CAS is no longer present), a “residual” effect emerges that is marked by different behavioral components. While the meanings of freezing and erratic swimming are likely context-dependent, and much up to debate, it could be argued that they represent “passive” or “risk assessment” responses in the case of freezing, and “active” or “escape” responses in the case of erratic swimming. Thus, as CAS is no longer present, the animal transitions from a “risky” to a “safety” behavioral state, as the absence of predators or partial predator stimuli are used as a positive prediction error. This is similar to what is observed with electrical stimulation of the PAG/GR in rats (Brandão et al., 2008). We have previously shown that serotonin differentially mediates these behaviors, phasically inhibiting responses during exposure and tonically facilitating responses after exposure, and suggested that it is the phasic activation of serotonin receptors after exposure that represent prediction errors (Lima-Maximino et al., 2020). The continued absence of these threat signals sets negative expectation values, proposed by Amo et al. (2014) to be encoded by raphe serotonergic neurons.

ARS-elicited behavioral effects have been explored in the literature with mixed results. While Ghisleni et al. (2012) found that a 90-min restraint protocol did not change bottom-dwelling after stress when animals were tested individually, a similar protocol showed marked increases in this variable (Assad et al., 2020) – albeit control animals in the latter experiment spent most of the session in the upper half of the tank instead of in the lower half. It is difficult to find other measures which are differentially affected by CAS vs. ARS, as physiological endpoints such as cortisol are increased by both (Ghisleni et al., 2012; Abreu et al., 2017). In the present experiments, ARS increased bottom-dwelling as well. Moreover, we observed a small component of erratic swimming, consistent with what is observed by Ghisleni et al. (2012). While increased bottom-dwelling was also observed as a pattern of post-exposure behavior after CAS in the present experiments and in Lima-Maximino et al. (2020), freezing was a major component of post-exposure behavior, suggesting that stressors do not produce similar behavioral effects.

### 4.2. Role of the 5-HT_2C_ receptor in CAS-elicited behavioral adjustments

The pattern of drug effects across behavioral experiments are complex, and summarized on Table 9. Both 5-HT_2C_ receptor agonists were able to block CAS-elicited behavioral adjustments *during* exposure, but the effects on behavior after exposure were less impressive, with agonists blocking the increased freezing. RS-102221, a 5-HT_2C_ receptor antagonist, blocked changes in erratic swimming and freezing during exposure, and blocked increases in geotaxis and freezing after exposure. These results suggest that the 5-HT_2C_ receptor has opposite roles in both stages, phasically inhibiting defensive responses to proximal threat and tonically facilitating responses to potential threat.

**Table 9.** Behavioral effects of 5-HT2C receptor agonists (MK-212 and WAY-161503) and antagonists (RS-102221) on behavior during and after CAS exposure, and after ARS in adult zebrafish.

| Experiment |  | Behavior |  |  |
| --- | --- | --- | --- | --- |
|  |  | Geotaxis | Erratic swimming | Freezing |
| During CAS exposure | CAS effect | ↑↑ | ↑↑ | ↑ |
|  | Agonist effect | Block | Partial block | Block |
|  | Antagonist effect | -- | Block | Partial block |
| After CAS exposure | CAS effect | ↑ | ↑ | ↑↑ |
|  | Agonist effect | -- | -- | Partial block |
|  | Antagonist effect | Block | -- | Partial Block |
| After ARS | ARS effect | ↑ | ↑ | ↑ |
|  | Agonist effect | -- | -- | Block |
|  | Antagonist effect | Partial block | -- | -- |

In the experimental design that was used in this experiment, behavior during exposure represents distal threat, as CAS acts as a partial predator stimulus that elicits behavior that decreases the possibility of a predator attack or detection by the predator (Smith, 1992).

However, if CAS is no longer present (as in the post-exposure stage of our design), that would signal a decrease in threat levels to potential threat, a situation in which trying to flee or hide is non-adaptive, but resuming normal behavior is also non-adaptive -- and therefore cautious, alert exploration is warranted (Maximino et al., 2019). Using Fanselow’s taxonomy, the situation in which CAS is present is more akin to “fear”, while the situation in which it is no longer present is more akin to “anxiety” (Perusini and Fanselow, 2015). The rodent literature suggests that 5-HT_2_ receptors participate in both fear and anxiety. Salchner and Singewald (2006) have shown that MK-212 potentiates escape responses to an airjet in rats, and the antagonist SB-242084 has been shown to produce anxiolytic-like effects in rats (Kennett et al., 1997; Martin et al., 2002). In the elevated T-maze, an apparatus which tests anxiety-like behavior (inhibitory avoidance) and fear-like behavior (one-way escape) in rodents, 5-HT_2C_ agonists facilitate, and antagonists impair, inhibitory avoidance, but only agonists facilitate one-way escape (Mora et al., 1997), suggesting tonic facilitation of responses to potential threat and phasic inhibition of responses to distal or proximal threat. Thus, a conserved role for this receptor appears to be related to phasic and tonic modulation of defensive responses.

We have previously found that serotonin participates in responses to CAS, acutely and phasically inhibiting responses to distal threat but phasically facilitating responses to potential threat (Lima-Maximino et al., 2020; Maximino et al., 2014). While it is not known whether serotonin levels vary during exposure (i.e., whether CAS elicits physiological, adaptive changes in serotonin levels), pharmacological experiments (Lima-Maximino et al., 2020) show that acute (phasic) increases in serotonin during exposure block behavioral effects of CAS, while decreasing serotonin levels with *para*-chlorophenylalanine did not produce effects during exposure. In the present experiments, 5-HT_2C_ receptor agonists blocked the effects of CAS during exposure, suggesting that during exposure phasic activation of the 5-HT_2C_ receptor is inhibitory. However, in the present experiments the antagonist RS-102221 also blocked effects on erratic swimming and (partially) freezing during exposure. In Nile tilapia, the 5-HT_2A/2C_ and α_2_-adrenoceptor antagonist mianserin blocked active components of the alarm reaction (dashing, bristling of dorsal fin spines), but not freezing, during exposure (Barreto, 2012). If erratic swimming and geotaxis are indeed the main components of the alarm reaction during exposure, the 5-HT_2C_ receptor could inhibit concurrent risk assessment (freezing) responses while facilitating active escape/avoidance responses (erratic swimming).

We have found that, after exposure, CAS increases serotonin levels in the extracellular fluid of the zebrafish brain (Maximino et al., 2014), an effect that can be mediated by serotonin transporters (Maximino et al., 2014) and/or inhibition of monoamine oxidase activity (Lima-Maximino et al., 2020). Regardless of the mechanism, increased serotonin levels *after* exposure are suggested to encode an aversive expectation value, switching behavior toward cautious exploration/risk assessment/anxiety when the aversive stimulus is no longer present (Lima-Maximino et al., 2020). With increased serotonin levels, receptor antagonists are expected to produce effects at this stage. This increase in serotonergic activity could activate 5-HT_2C_ and other serotonergic receptors to inhibit ongoing defensive responses to proximal threat (e.g., freezing) and initiate programs of alertness and cautious exploration/risk assessment, including geotaxis and erratic swimming (which are main components of the response during exposure). However, while in the present experiments RS-102221, a 5-HT_2C_ receptor antagonist, blocked increases in erratic swimming and geotaxis observed after exposure, 5-HT_2C_ receptor agonists blocked freezing observed after exposure, contrary to what would be expected if that was true. Moreover, 5-HT_2C_ receptors have been shown to inhibit 5-HT neuronal discharge in anesthetized rats (Queree et al., 2009) through a regulatory loop mediated by GABAergic neurons in the raphe, habenula, and/or frontal cortex (Sharp et al., 2007). Thus, the effects of receptor agonists in this stage could represent the over-activation of this regulatory loop in conditions of elevated serotonin levels. Moreover, other receptors can be implicated; indeed, in zebrafish, both the 5-HT_2_ receptor antagonist methysergide and the 5-HT1A receptor antagonist WAY-100635 increased freezing and bottom-dwelling during CAS exposure and during a recovery period (Nathan et al., 2015), suggesting that other receptors from the 5-HT_2_ family or the 5-HT_1A_ receptor (but see Maximino et al., 2014) control freezing behavior after exposure. Thus, serotonergic signals emerging after exposure, representing negative expectation values (Lima-Maximino et al., 2020), would modulate behavior through the 5-HT_2C_ receptor (inhibiting active responses that would emerge in distal threat) and other receptors (facilitating freezing/risk assessment responses that emerge in potential threat).

However, at least for the case of the 5-HT_2C_ receptor, the phasic signals do not appear to represent all aversive values, as no effect of MK-212 was found for most ARS-elicited responses, with the exception of freezing. Moreover, RS-102221 blocked the effects of ARS on geotaxis. In rats, acute restraint stress increases serotonin release in the central amygdala, an effect that is mediated by corticotropin-releasing factor receptors (Mo et al., 2008). In mce, restraint stress for 45 min. increased serotonin turnover in the hippocampus, an effect blocked by the 5-HT2_B/2C_ receptor agonist RO 60-0175 (Mongeau et al, 2010). In Nile tilapia, acute restraint stress (20 min) increased serotonin turnover in the dorsolateral and dorsomedial telencephali (Silva et al., 2015), homologous to the hippocampus and frontotemporal amygdala complex, respectively. Interestingly, the 5-HT_2C_ receptor antagonist SB 242084 blocked the inhibitory effect of restraint stress on social interactions in mice (Mongeau et al., 2010). Thus, the effects of RS-102221 on geotaxis could be mediated by blockade of 5-HT_2C_ receptors in the forebrain, while the effects of MK-212 would be more likely to involve activation of the regulatory loop. Our results suggest that the 5-HT_2C_ receptor is at least partially responsible for both effects, acting as a “switch” between behavioral modes.

One potential limitation of behavioral pharmacological work in zebrafish is the relative divergence between 5-HT_2C_-like receptors in teleosts and mammals, which makes it difficult to ascertain whether drugs which act as agonists and antagonists at mammalian 5-HT_2C_ receptors will target zebrafish receptors. Using sequence alignment, it was found that residues that, in human receptors, are crucial for ligand binding (and selectivity with the 5-HT_2A_ receptor, for example; Canal et al., 2011; Córdova-Sintjago et al., 2012) are conserved in both zebrafish proteins (HTR2Cl1 and HTR2Cl2). This suggests that ligand affinity is conserved for these proteins, which strengthens the hypothesis that MK-212, WAY-161503, and RS-102221 targeted 5-HT_2C_ receptors in the behavioral experiments. Differences were found in residues near the conserved binding sites, however; even in mammalian receptors, the role of these residues in ligand affinity and/or efficacy is unknown.

### 4.3. Limitations and translational potential

While the present work presents results on zebrafish behavioral pharmacology inspired on results from mammals, which can potentially underscore conserved mechanisms for responses to threat across vertebrates (cf. Gerlai, 2014, for a discussion on how using fish as well as mammals can decrease uncertainty regarding translational relevance and evolutionary conservation of mechanisms), there are limitations in using zebrafish as experimental models in neuroscience. One of them is that, due to the teleost genome duplication event, zebrafish present two copies of the gene that codes the 5-HT_2C_ receptor (Schneider et al., 2012), and therefore it is currently not possible to know which protein (HTR2CL1 or HTR2Cl2) was targeted by the drugs used in the present experiments. Importantly, both proteins had conserved residues at sites known to affect ligand affinity in the human 5-HT_2C_ receptor, and therefore both are likely to have produced the effects. Moreover, while we propose that alarm reactions and post-exposure behavior can be exploited to better understand panic disorder (Silva et al., 2020), the paradigm that was used represents the normal, adaptive behavior of zebrafish under threatening circumstances, and therefore cannot yet be used to generate novel hypotheses about a pathological organism (cf. Maximino and van der Staay, 2019, for a discussion on the limitations of using behavioral assays in experimental psychopathology).

Another important limitation of our results is that sex differences were not considered throughout the experiment. The field of sex differences in zebrafish research is barely emerging (Genario et al., 2020), due to difficulties in ascertaining sex only through external morphology in common (commercial) zebrafish phenotypes; as a result, we did not separate animals per sex in our experiments. Future experiments will consider the role of sex in these responses, considering that sex differences were observed in fear-like behavior in rodents (e.g., Petrovich and Lougee, 2011).

Despite these limitations, the work presented has translational potential in understanding fear- and anxiety-like states (cf. Silva et al., 2020, for a discussion on the use of alarm reactions to study fear in zebrafish). In general, our results are the first to determine a specific role of 5-HT_2C_ receptors in zebrafish behavior, and add to the small literature on the role of this receptor in mammals as well. Further work is needed to understand whether this receptor interacts with other serotonin receptors known to be involved in defensive behavior in this species (Herculano and Maximino, 2014) and if the effects during exposure are related to changes in serotonin levels. Moreover, given the importance of these phenotypes to understanding defensive behavior (Silva et al., 2020), further work will clarify the usefulness of this pharmacological profile in modeling fear and anxiety.

## Significance statement

While the neurotransmitter serotonin has been associated with mental disorders, we currently do not know the mechanisms through which serotonin influences responses to threat. This is relevant to future studies on anxiety disorders, for example. Here we report that, in zebrafish, the 5-HT_2C_ receptor “pushes the break” on (fear-like) responses to threat that is actually present, and keeps on facilitating (anxiety-like) responses to threat that is only implied. Thus, this receptor appears to act as a “switch” between two modes that facilitate escape (“fear”) or cautious exploration (“anxiety”).

## Acknowledgments

This work was supported by a Conselho Nacional de Desenvolvimento Científico e Tecnológico (CNPq) grant (#400726/2016-5). CM is the recipient of a CNPq productivity grant (PQ2, #302998/2019-5). The funders had no influence on the study design, the collection, analysis and interpretation of data, as well as writing and submission of this manuscript.

## Authors’ contributions

*Rhayra Xavier do Carmo Silva:* Conceptualization, Methodology, Investigation, Data curation, Formal analysis, Visualization, Writing – original draft, Writing – review & editing; *Bianca Gomes do Nascimento:* Investigation, Data curation, Formal analysis; *Gabriela Cristini Vidal Gomes:* Investigation, Data curation, Formal analysis; *Nadyme Assad Holanda da Silva:* Investigation, Data curation; *Jéssica Souza Pinheiro:* Investigation, Data curation; *Suianny Nayara da Silva Chaves:* Investigation, Data curation, Formal analysis, Visualization; *Ana Flávia Nogueira Pimentel:* Investigation, Data curation, Formal analysis; *Bruna Patrícia Dutra Costa:* Investigation, Data curation, Formal analysis; *Anderson Manoel Herculano:* Conceptualization, Supervision, Resources, Validation; *Monica Lima-Maximino:* Conceptualization, Validation, Writing – original draft; *Caio Maximino:* Conceptualization, Formal analysis, Funding acquisition, Methodology, Project Administration, Resources, Software, Supervision, Validation, Visualization, Writing – original draft, Writing – review & editing.

## Data accessibility

Data and analysis scripts for this work can be found at a GitHub repository (https://github.com/lanec-unifesspa/5-HT-CAS/tree/master/data/5HT2C/; doi: 10.5281/zenodo.4437275).

## References

1. Alves, S.H., Pinheiro, G., Motta, V., Landeira-Fernandez, J., Cruz, A.P.M., 2004. Anxiogenic effects in the rat elevated plus-maze of 5-HT2C agonists into ventral but not dorsal hippocampus. Behav. Pharmacol. 15, 37–43.

2. Amo, R., Fredes, F., Kinoshita, M., Aoki, R., Aizawa, H., Agetsuma, M., Aoki, T., Shiraki, T., Kakinuma, H., Matsuda, M., Yamazaki, M., Takahoko, M., Tsuboi, T., Higashijima, S., Miyasaka, N., Koide, T., Yabuki, Y., Yoshihara, Y., Fukai, T., Okamoto, H., 2014. The Habenulo-Raphe Serotonergic Circuit Encodes an Aversive Expectation Value Essential for Adaptive Active Avoidance of Danger. Neuron 84, 1034–1048. https://doi.org/10.1016/j.neuron.2014.10.035

3. Assad, N., Luz, W.L., Santos-Silva, M., Carvalho, T., Moraes, S., Picanço-Diniz, D.L.W., Bahia, C.P., Oliveira Batista, E. de J., da Conceição Passos, A., Oliveira, K.R.H.M., Herculano, A.M., 2020. Acute Restraint Stress Evokes Anxiety-Like Behavior Mediated by Telencephalic Inactivation and GabAergic Dysfunction in Zebrafish Brains. Sci. Rep. 10, 5551. https://doi.org/10.1038/s41598-020-62077-w

4. Ballesteros, J. A., Jensen, A. D., Liapakis, G., Rasmussen, S. G., Shi, L., Gether, U., Javitch, J. A., 2001. Activation of the beta 2-adrenergic receptor involves disruption of an ionic lock between the cytoplasmic ends of transmembrane segments 3 and 6. J. Biol. Chem. 276, 29171–29177. https://doi.org/10.1074/jbc.M103747200

5. Barreto, R.E., 2012. Mianserin affects alarm reaction to conspecific chemical alarm cues in Nile tilapia. Fish Physiol. Biochem. 43, 193–201. https://doi.org/10.1007/s10695-016-0279-2

6. Bencan, Z., Sledge, D., Levin, E.D., 2009. Buspirone, chlordiazepoxide and diazepam effects in a zebrafish model of anxiety. Pharmacol. Biochem. Behav. 94, 75–80. https://doi.org/10.1016/j.pbb.2009.07.009

7. Bonapersona, V., Hoijtink, H., Relacs Sarabdjitsingh, R.A., Joëls, M., 2019. RePAIR: a power solution to animal experimentation. bioRxiv 864652. https://doi.org/10.1101/864652

8. Bonhaus, D.W., Weinhardt, K.K., Taylor, M., DeSouza, A., McNeeley, P.M., Szczepanski, K., Fontana, D.J., Trinh, J., Rocha, C.L., Dawson, M.W., Flippin, L.A., Eglen, R.M., 1997. RS-102221: a novel high affinity and selective, 5-HT2C receptor antagonist. Neuropharmacology 36, 621–629. https://doi.org/10.1016/s0028-3908(97)00049-x

9. Brandão, M.L., Zanoveli, J.M., Ruiz-Martinez, R.C., Oliveira, L.C., Landeira-Fernandez, J., 2008. Different patterns of freezing behavior organized in the periaqueductal gray of rats: Association with different types of anxiety. Behav. Brain Res. 188, 1–13. https://doi.org/10.1016/j.bbr.2007.10.018

10. Canal, C. E., Cordova-Sintjago, T. C., Villa, N. Y., Fang, L.-J., & Booth, R. G. (2011). Drug discovery targeting human 5-HT2C receptors: Residues S3.36 and Y7.43 impact ligand—Binding pocket structure via hydrogen bond formation. Eur. J. Pharmacol., 673, 1–12. https://doi.org/10.1016/j.ejphar.2011.10.006

11. Castilho, V., Macedo, C., Brandão, M., 2002. Role of benzodiazepine and serotonergic mechanisms in conditioned freezing and antinociception using electrical stimulation of the dorsal periaqueductal gray as unconditioned stimulus in rats. Psychopharmacology (Berl.) 165, 77–85. https://doi.org/10.1007/s00213-002-1246-4

12. Castilho, V.M., Brandão, M.L., 2001. Conditioned antinociception and freezing using electrical stimulation of the dorsal periaqueductal gray or inferior colliculus as unconditioned stimulus are differentially regulated by 5-HT2A receptors in rats. Psychopharmacology (Berl.) 155, 154–162. https://doi.org/10.1007/s002130100697

13. Coimbra, N.C., Brandão, M.L., 1997. Effects of 5-HT2 receptors blockade on fear-induced analgesia elicited by electrical stimulation of the deep layers of the superior colliculus and dorsal periaqueductal gray. Behav. Brain Res. 87, 97–103. https://doi.org/10.1016/S0166-4328(96)02267-X

14. Conselho Nacional de Controle de Experimentação Animal - CONCEA, 2017. Diretriz brasileira para o cuidado e a utilização de animais para fins científicos e didáticos - DBCA. Anexo I. Peixes mantidos em instalações de instituições de ensino ou pesquisa científica, Resolução Normativa CONCEA no 34.

15. Córdova-Sintjago, T., Sakhuja, R., Kondabolu, K., Canal, C. E., Booth, R. G., 2012. Molecular determinants of ligand binding at serotonin 5-HT_2A_ and 5-HT_2C_ GPCRs: Experimental affinity results analyzed by molecular modeling and ligand docking studies. Int. J. Quantum Chemistry, 112, 3807–3814.

16. Córdova-Sintjago, T., Sakhuja, R., Kondabolu, K., Canal, C. E., & Booth, R. G. (2012). Molecular determinants for ligand binding at serotonin 5-HT2Aand 5-HT2CGPCRs: Experimental affinity results analyzed by molecular modeling and ligand docking studies. International Journal of Quantum Chemistry, 112(24), 3807–3814. doi:10.1002/qua.24237

17. Cornélio, A.M., Nunes-de-Souza, R.L., 2007. Anxiogenic-like effects of mCPP microinfusions into the amygdala (but not dorsal or ventral hippocampus) in mice exposed to elevated plus-maze. Behav. Brain Res. 178, 82–89. https://doi.org/10.1016/j.bbr.2006.12.003

18. de Mello Cruz, A.P., Pinheiro, G., Alves, S.H., Ferreira, G., Mendes, M., Faria, L., Macedo, C.E., Landeira-Fernandez, J., 2005. Behavioral effects of systemically administered MK-212 are prevented by ritanserin microinfusion into the basolateral amygdala of rats exposed to the elevated plus-maze. Psychopharmacology (Berl.) 182, 345–354. https://doi.org/10.1007/s00213-005-0108-2

19. de Paula, B.B., Leite-Panissi, C.R.A., 2016. Distinct effect of 5-HT1A and 5-HT2A receptors in the medial nucleus of the amygdala on tonic immobility behavior. Brain Res. 1643, 152– 158. https://doi.org/10.1016/j.brainres.2016.04.073

20. do Carmo Silva, R.X., Lima-Maximino, M.G., Maximino, C., 2018a. The aversive brain system of teleosts: Implications for neuroscience and biological psychiatry. Neurosci. Biobehav. Rev. 95, 123–135. https://doi.org/10.1016/j.neubiorev.2018.10.001

21. do Carmo Silva, R.X., Rocha, S.P., Lima-Maximino, M.G., Maximino, C., 2018b. Extraction of alarm substance in zebrafish. protocols.io. https://doi.org/10.17504/protocols.io.tr3em8n

22. Genario, R., Abreu, M. S., Giacomini, A. C. V. V., Demin, K. A., Kalueff, A. V. Sex differences in behavior and neuropharmacology of zebrafish. Eur. J. Neurosci. 51, 2586–2603. https://doi.org/10.1111/ejn.14438

23. Gerlai. R., 2014. Fish in behavior research: Unique tools with a great promise! J. Neurosci. Methods 234, 54–58. https://doi.or/g10.1016/j.jneumeth.2014.04.015

24. Ghisleni, G., Capiotti, K.M., Da Silva, R.S., Oses, J.P., Piato, Â.L., Soares, V., Bogo, M.R., Bonan, C.D., 2012. The role of CRH in behavioral responses to acute restraint stress in zebrafish. Prog. Neuropsychopharmacol. Biol. Psychiatry 36, 176–182. https://doi.org/10.1016/j.pnpbp.2011.08.016

25. Gomes, F., Greidinger, M., Salviano, M., Couto, K.C., Scaperlli, G.F., Alves, S.H. de S., Cruz, A.P. de M., 2010. Antidepressant- and anxiogenic-like effects of acute 5-HT2C receptor activation in rats exposed to the forced swim test and elevated plus maze. Psychol. & Neurosci. 3, 245–249. https://doi.org/10.3922/j.psns.2010.2.014

26. Goodwin, N., Karp, N. A., Blackledge, S., Clark, B., Keeble, R., Kovacs, C., Murray, K. N., Price, M., Thompson, P., Bussell J., 2016. Standardized welfare terms for the zebrafish community. Zebrafish 13, Suppl. 1, S164-S168. https://doi.org/10.1089/zeb.2016.1248

27. Graeff, F.G., 2004. Serotonin, the periaqueductal gray and panic. Neurosci. Biobehav. Rev. 28, 239–259. https://doi.org/10.1016/j.neubiorev.2003.12.004

28. Graeff, F.G., Brandão, M.L., Audi, E.A., Schütz, M.T.B., 1986. Modulation of the brain aversive system by GABAergic and serotonergic mechanisms. Behav. Brain Res. 22, 173–180. https://doi.org/10.1016/0166-4328(86)90038-0

29. Herculano, A.M., Maximino, C., 2014. Serotonergic modulation of zebrafish behavior: Towards a paradox. Prog. Neuropsychopharmacol. Biol. Psychiatry, Special Issue: Zebrafish models of brain disorders 55, 50–66. https://doi.org/10.1016/j.pnpbp.2014.03.008

30. Höglund, E., Weltzien, F.-A., Schjolden, J., Winberg, S., Ursin, H., Døving, K.B., 2005. Avoidance behavior and brain monoamines in fish. Brain Res. 1032, 104–110. https://doi.org/10.1016/j.brainres.2004.10.050

31. Kennett, G.A., Wood, M.D., Bright, F., Trail, B., Riley, G., Holland, V., Avenell, K.Y., Stean, T., Upton, N., Bromidge, S., Forbes, I.T., Brown, A.M., Middlemiss, D.N., Blackburn, T.P., 1997. SB 242084, a Selective and Brain Penetrant 5-HT2C Receptor Antagonist. Neuropharmacology 36, 609–620. https://doi.org/10.1016/S0028-3908(97)00038-5

32. Kinkel, M.D., Eames, S.C., Philipson, L.H., Prince, V.E., 2010. Intraperitoneal Injection into Adult Zebrafish. JoVE J. Vis. Exp. e2126. https://doi.org/10.3791/2126

33. Knight, A.R., Misra, A., Quirk, K., Benwell, K., Revell, D., Kennett, G., Bickerdike, M., 2004. Pharmacological characterisation of the agonist radioligand binding site of 5-HT2A, 5-HT2B and 5-HT2C receptors. Naunyn. Schmiedebergs Arch. Pharmacol. 370, 114–123. https://doi.org/10.1007/s00210-004-0951-4

34. Kuznetsova, E.G., Amstislavskaya, T.G., Shefer, E.A., Popova, N.K., 2006. Effect of 5-HT2C receptor antagonist RS 102221 on mouse behavior. Bull. Exp. Biol. Med. 142, 76–79. https://doi.org/10.1007/s10517-006-0296-8

35. Lima-Maximino, M., Pyterson, M.P., Silva, R.X. do C., Gomes, G.C.V., Rocha, S.P., Herculano, A.M., Rosemberg, D.B., Maximino, C., 2020. Phasic and tonic serotonin modulate alarm reactions and post-exposure behavior in zebrafish. J. Neurochem. 153, 495–509. https://doi.org/10.1111/jnc.14978

36. Macedo, C.E., Martinez, R.C.R., Albrechet-Souza, L., Molina, V.A., Brandão, M.L., 2007. 5-HT2- and D1-mechanisms of the basolateral nucleus of the amygdala enhance conditioned fear and impair unconditioned fear. Behav. Brain Res. 177, 100–108. https://doi.org/10.1016/j.bbr.2006.10.031

37. Martin, J.R., Ballard, T.M., Higgins, G.A., 2002. Influence of the 5-HT2C receptor antagonist, SB-242084, in tests of anxiety. Pharmacol. Biochem. Behav., Serotonin: Pharmacology, Biochemistry & Behaviour 71, 615–625. https://doi.org/10.1016/S0091-3057(01)00713-4

38. Mathuru, A.S., Kibat, C., Cheong, W.F., Shui, G., Wenk, M.R., Friedrich, R.W., Jesuthasan, S., 2012. Chondroitin Fragments Are Odorants that Trigger Fear Behavior in Fish. Curr. Biol. 22, 538–544. https://doi.org/10.1016/j.cub.2012.01.061

39. Matthews, M., Varga, Z.M., 2012. Anesthesia and Euthanasia in Zebrafish. ILAR J. 53, 192– 204. https://doi.org/10.1093/ilar.53.2.192

40. Maximino, C., van der Staay, J., 2019. Behavioral models in psychopathology: Epistemic and semantic considerations. Behav. Brain Funct. 15, 1. https://doi.org/10.1186/s12993-019-0152-4

41. Maximino, C., do Carmo Silva, R.X., Campos, K. dos S., de Oliveira, J.S., Rocha, S.P., Pyterson, M.P., Souza, D.P. dos S., Feitosa, L.M., Ikeda, S.R., Pimentel, A.F.N., Ramos, P.N.F., Costa, B.P.D., Herculano, A.M., Rosemberg, D.B., Siqueira-Silva, D.H., Lima-Maximino, M., 2019. Sensory ecology of ostariophysan alarm substances. J. Fish Biol. 95, 274–286. https://doi.org/10.1111/jfb.13844

42. Maximino, C., Lima, M.G., Costa, C.C., Guedes, I.M.L., Herculano, A.M., 2014. Fluoxetine and WAY 100,635 dissociate increases in scototaxis and analgesia induced by conspecific alarm substance in zebrafish (*Danio rerio* Hamilton 1822). Pharmacol. Biochem. Behav. 124, 425–433. https://doi.org/10.1016/j.pbb.2014.07.003

43. Mo, B., Feng, N., Renner, K., Forster, G., 2008. Mo, B., Feng, N., Renner, K., & Forster, G. (2008). Restraint stress increases serotonin release in the central nucleus of the amygdala via activation of corticotropin-releasing factor receptors. Brain Res. Bul. 76, 493–498. https://doi.org/10.1016/j.brainresbull.2008.02.011

44. Mora, P.O., Netto, C.F., Graeff, F.G., 1997. Role of 5-HT2A and 5-HT2C Receptor Subtypes in the Two Types of Fear Generated by the Elevated T-Maze. Pharmacol. Biochem. Behav. 58, 1051–1057. https://doi.org/10.1016/S0091-3057(97)00057-9

45. Nathan, F.M., Ogawa, S., Parhar, I.S., 2015. Kisspeptin1 modulates odorant-evoked fear response via two serotonin receptor subtypes (5-HT1A and 5-HT2) in zebrafish. J. Neurochem. 133, 870–878. https://doi.org/10.1111/jnc.13105

46. Nunes-de-Souza, V., Nunes-de-Souza, R.L., Rodgers, R.J., Canto-de-Souza, A., 2008. 5-HT2 receptor activation in the midbrain periaqueductal grey (PAG) reduces anxiety-like behaviour in mice. Behav. Brain Res. 187, 72–79. https://doi.org/10.1016/j.bbr.2007.08.030

47. Oliveira, L.C., Broiz, A.C., de Macedo, C.E., Landeira-Fernandez, J., Brandão, M.L., 2007. 5-HT2 receptor mechanisms of the dorsal periaqueductal gray in the conditioned and unconditioned fear in rats. Psychopharmacology (Berl.) 191, 253–262. https://doi.org/10.1007/s00213-006-0653-3

48. Parra, K.V., Adrian, J.C., Gerlai, R., 2009. The synthetic substance hypoxanthine 3-N-oxide elicits alarm reactions in zebrafish (Danio rerio). Behav. Brain Res. 205, 336–341. https://doi.org/10.1016/j.bbr.2009.06.037

49. Perusini, J.N., Fanselow, M.S., 2015. Neurobehavioral perspectives on the distinction between fear and anxiety. Learn. Mem. 22, 417–425. https://doi.org/10.1101/lm.039180.115

50. Petrovich, G. D., Lougee, M. A., 2011. Sex differences in fear-induced feeding cessation: Prolonged effect in female rats. Physiol. Behav. 104, 996–1101. https://doi.org/10.1016/j.physbeh.2011.06.020

51. Piato, A.L., Rosemberg, D.B., Capiotti, K.M., Siebel, A.M., Herrmann, A.P., Ghisleni, G., Vianna, M.R., Bogo, M.R., Lara, D.R., Bonan, C.D., 2011. Acute Restraint Stress in Zebrafish: Behavioral Parameters and Purinergic Signaling. Neurochem. Res. 36, 1876. https://doi.org/10.1007/s11064-011-0509-z

52. Quadros, V.A., Costa, F.V., Canzian, J., Nogueira, C.W., Rosemberg, D.B., 2018. Modulatory role of conspecific alarm substance on aggression and brain monoamine oxidase activity in two zebrafish populations. Prog. Neuropsychopharmacol. Biol. Psychiatry 86, 322– 330. https://doi.org/10.1016/j.pnpbp.2018.03.018

53. Quadros, V.A., Silveira, A., Giuliani, G.S., Didonet, F., Silveira, A.S., Nunes, M.E., Silva, T.O., Loro, V.L., Rosemberg, D.B., 2016. Strain- and context-dependent behavioural responses of acute alarm substance exposure in zebrafish. Behav. Processes 122, 1– 11. https://doi.org/10.1016/j.beproc.2015.10.014

54. Queree, P., Peters, S., Sharp, T., 2009. Further pharmacological characterization of 5-HT(2C) receptor agonist-induced inhibition of 5-HT neuronal activity in the dorsal raphe nucleus in vivo. Br. J. Pharmacol., 158, 1477–1485. https://doi.org/10.1111/j.1476-5381.2009.00406.x

55. Rosenzweig-Lipson, S., Zhang, J., Mazandarani, H., Harrison, B.L., Sabb, A., Sabalski, J., Stack, G., Welmaker, G., Barrett, J.E., Dunlop, J., 2006. Antiobesity-like effects of the 5-HT2C receptor agonist WAY-161503. Brain Res. 1073–1074, 240–251. https://doi.org/10.1016/j.brainres.2005.12.052

56. Salchner, P., Singewald, N., 2006. 5-HT receptor subtypes involved in the anxiogenic-like action and associated Fos response of acute fluoxetine treatment in rats. Psychopharmacology (Berl.) 185, 282–288. https://doi.org/10.1007/s00213-005-0247-5

57. Schneider, H., Fritzky, L., Williams, J., Heumann, C., Yochum, M., Pattar, K., Noppert, G., Mock, V., Hawley, E., 2012. Cloning and expression of a zebrafish 5-HT_2C_ receptor gene. Gene 502, 108–117. https://doi.org/10.1016/j.gene.2012.03.070

58. Sharp, T., Boothman, L., Raley, J., Queree, P., 2007. Important messages in the ‘post’: Recent discoveries in 5-HT neurone feedback control. Trends Pharmacol. Sci., 28, 629–635. https://doi.org/10.1016/j.tips.2007.10.009

59. Silva, P.I.M., Martins, C.I.M., Khan, U.W., Gjøen, H.M., Øverli, Ø., Höglund, E., 2015. Stress and fear responses in the teleost pallium. Physiol. Behav. 141, 17–22. https://doi.org/10.1016/j.physbeh.2014.12.020

60. Silva, R.X. do C., Rocha, S.P., Herculano, A.M., Lima-Maximino, M.G., Maximino, C., 2020. Animal models for panic disorder. Psychol. Neurosci. 13, 1–18. https://doi.org/10.1037/pne0000177

61. Smith, R.J.F., 1992. Alarm signals in fishes. Rev. Fish Biol. Fish. 2, 33–63. https://doi.org/10.1007/BF00042916

62. Sourbron, J., Schneider, H., Kecskés, A., Liu, Y., Buening, E.M., Lagae, L., Smolders, I., de Witte, P., 2016. Serotonergic Modulation as Effective Treatment for Dravet Syndrome in a Zebrafish Mutant Model. ACS Chem. Neurosci. 7, 588–598. https://doi.org/10.1021/acschemneuro.5b00342

63. Speedie, N., Gerlai, R., 2008. Alarm substance induced behavioral responses in zebrafish (Danio rerio). Behav. Brain Res. 188, 168–177. https://doi.org/10.1016/j.bbr.2007.10.031

64. von Frisch, K., 1938. Zur Psychologie des Fisch-Schwarmes. Naturwissenschaften 26, 601– 606. https://doi.org/10.1007/BF01590598

65. Wolf, K., 1963. Physiological Salines for Fresh-Water Teleosts. Progress. Fish-Cult. 25, 135– 140. https://doi.org/10.1577/1548-8659(1963)25[135:PSFFT]2.0.CO;2

